# Finding the E-Channel Proton Loading Sites by Calculating the Ensemble of Protonation Microstates

**DOI:** 10.1101/2024.08.14.608010

**Authors:** Md. Raihan Uddin, Umesh Khaniya, Chitrak Gupta, Junjun Mao, Gehan Ranepura, Rongmei Wei, Jose Ortiz-Soto, Abhishek Singharoy, M.R. Gunner

## Abstract

The aerobic electron transfer chain builds a proton gradient by proton coupled electron transfer reactions through multiple proteins. Complex I is the first enzyme in this chain. On transfer of two electrons from NADH to quinone four protons are pumped from the N- (negative, higher pH) to the P- (positive, lower) side. Protons move pthrough three linear antiporter paths, with a few amino acids and waters providing the route and the E-channel, which is a complex of competing paths, with clusters of interconnected protonatable residues.

Proton loading sites (PLS) are residues that transiently bind protons as they are transported from N- to P-compartments. PLS can be individual residues or extended clusters. The program MCCE uses Monte Carlos sampling to analyze the E-channel proton binding in equilibrium with individual Molecular Dynamics snapshots from trajectories of *Thrmus thermuphillus* Complex I in the apo, quinone or quinol bound states. At pH 7, the five E-channel subunits (Nqo4, Nqo7, Nqo8, Nqo10, and Nqo11) have accepted >25,000 protonation microstates, each with different residues protonated. The microstate explosion is tamed by analyzing interconnected clusters of residues along the proton transfer paths. A proton is bound and released to five coupled residues in a cluster on the protein N-side and to six coupled residues in the protein center. Loaded microstates bind protons to sites closer to the P-side in the forward pumping direction. MCCE microstate analysis shows the strong coupling of proton binding amongst individual residues in the two PLS clusters.

## 1. Introduction

The enzymes of the aerobic bacterial and mitochondrial electron transfer chains are multi-subunit protein assemblies, denoted as Complexes I–IV. Complex I, the first and largest enzyme, connects the soluble Krebs cycle reactions to those of the transmembrane electron transfer chain. It extends ∼200 Å within the membrane with an ∼150Å periplasmic arm. Proton pumps, such as Complex I, store energy via chemiosmotic coupling [1]. Protons are moved to the lower pH, more positive (P-side) compartment driven by redox reactions, moving electrons downhill to sites at more positive potential. The proton pumps establish an electrochemical proton motive force ranging from 100 to 200 mV across the membrane [1], which drives the F_1_F_0_-ATP synthase [2, 3]. In mitochondria, Complex I contribute approximately 40% of the proton flux that builds the gradient [1–5].

The energy released during the transfer of two-electrons from NADH to ubiquinone in the periplasmic domain is coupled to the translocation of four protons across the membrane domains [6]. The overall reaction is:

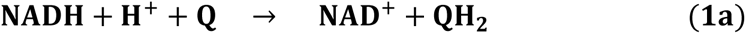

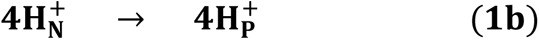

There are four proposed proton transfer paths: the E-channel, which is closest to the peripheral subunit where electron transfer takes place and three homologous antiporter domains [6]. It has been proposed that each antiporter subunit transports one proton [7, 8] with the fourth proton traveling through the E-channel [8–10], though the number and identity of pumping pathways are by no means settled [6, 9, 11, 12].

Protons move across the membrane embedded proton pumps such as Complex I [13], bacteriorhodopsin [14], and Cytochrome c Oxidase (CcO) [15, 16]. Conceptually, three elements are needed to drive protons uphill from the membrane N- to P-side [13, 16]: paths, loading sites (PLS), and gates. Hydrogen-bonded paths permit proton transfer via the Grotthuss Mechanism [17, 18]. The three antiporter paths follow a linear pattern, employing a few amino acids and water molecule molecules to create a route for protons. In contrast, the E-channel is complex, comprising competing paths. The E-channel has been modeled using Molecular Dynamics (MD) simulations [7, 19, 20], and by MCCE (MultiConformation Continuum Electrostatics) Monte Carlo (MC) sampling [10].

Protons are temporarily held within a proton loading site (PLS), where they are bound and then released as part of the reaction cycle. PLS residues lie along the proton transfer path and can be a single amino acid [21] or clusters of residues [10]. During the reaction sequence the proton affinity of the PLS rises to bind a proton and then is reduced to release it [18, 21, 22]. In CcO changes in hydrogen bonding patterns [21, 23] or entry of water molecule [24] lead to changes in proton affinity of different PLS of over 5 kcal/mol. The electrochemical gradient across the membrane encourages energy dissipating proton transfer, from the P- to the N-side. Therefore, during proton loading, the PLS must connect to the N-side and to the P-side during proton release, with gates blocking proton transfers in the wrong direction.

Crystal structures and cryo-EM structures [9, 12, 25–35] of Complex I have laid the foundation for in-depth investigations into proton pumping paths and the energy conserving pumping mechanism. Specific buried ionizable residues (Asp, Glu, and Lys) have been identified as possible PLS, members of proton transfer paths or as stabilizing needed internal hydration [6, 7, 20]. The importance of many of these residues has been confirmed by site-directed mutations [36–38].

Multiple computational methods have been applied to find the proton transfer paths and the PLS, to determine what triggers the protons to bind and release. Simulations have often focused on the linear antiporter channels [19, 39–42]. MD simulations [19, 41–43] have been instrumental in delineating hydrogen-bonded paths, often using trajectories with different fixed buried residue protonation states and cofactor redox states. Quantum Mechanics/Molecular Mechanics (QM/MM) methods have further probed local proton transfer events [19, 41, 42]. MD simulations involving the membrane domain of *E. coli* Complex I show that changes in protonation states of key residues can impact the hydration of channels supporting proton transfer [19, 43]. Generally, the calculated paths through the antiporter subunits align with those predicted based on the location of charged and polar residues in the crystal structure [41, 42].

The antiporter subunits are related to the Mrp Na/H antiporters [44] and the redox active periplasmic subunit is related to soluble hydrogenases [45–47] (Fig. 1). However, the E-channel is unique to Complex I. It is made up of five subunits, including Nqo4 that makes up the quinone binding site in the periplasmic arm, and the integral membrane Nqo7, 8, 10 and 11. (*T. thermophilus* naming used here See Table SI.1 for naming conventions for Complex I from other organisms) (Fig. 1b). The E-channel does not form a linear proton transfer path, rather there are many residues that form competing paths [10].

**Fig. 1.**
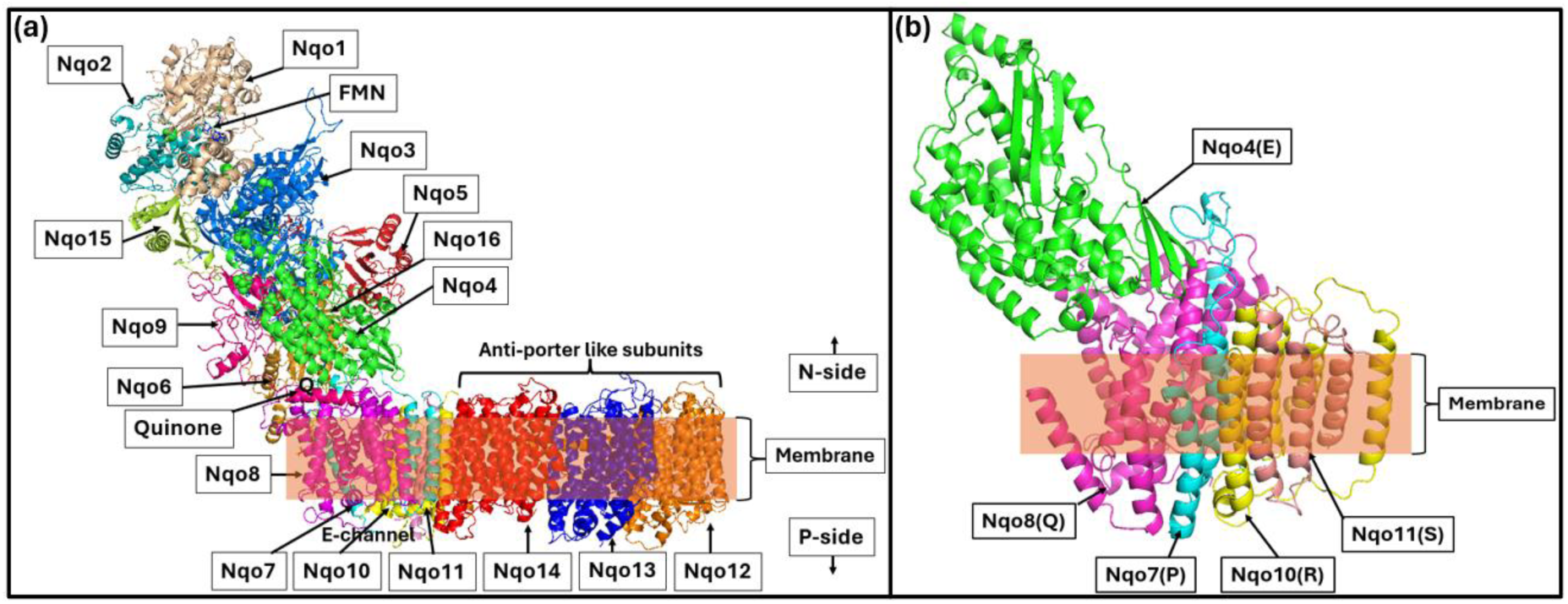
(a) MD frame of complete Complex I without MD membrane, ions, iron-sulfur clusters, and water molecules shown. (b) Five subunits extracted from the complete Complex I are used in MCCE simulation. Nqo numbers give the subunit identifiers used for the *T. thermophilus* Complex I and letters (E, P, Q, R, K) are the chain ID in the PDB file used (PDB ID: 4HEA).

PLS are defined here as being along the proton transfer path and being in different protonation states in different reaction intermediates. Thus, to identify them via computation, we need structures in different reaction states and a procedure to find the high probability protonation states. In Complex I quinone binding, reduction and release are assumed to trigger proton pumping [13, 38, 47–50], so apo, quinone- and quinol-bound states have been trapped experimentally [6, 51, 52] and modeled with MD [19, 51, 53, 54]. The connectivity of proton transfer paths has been evaluated in Complex I structures prepared in different states [51, 53]. However, MD and QM/MM must be carried out with fixed protonation states. Thus, the simulations are required to make educated guesses as to which residues bind protons, and these choices are unchanged through the simulation.

The MCCE program has been used to trace proton transfer paths in different proteins [10, 21, 22, 55, 56]. The program uses MC sampling to find the Boltzmann distribution of side chain position and protonation states given a fixed protein backbone [57]. MCCE finds the hydrogen bonds and protonation states that are at equilibrium with the input structure. This stochastic method allows us to characterize paths through a complex network. MCCE cannot make large conformational changes but brings the protonation states into equilibrium with the structure. Multiple snapshots from MD trajectories, each run with fixed protonation, allow for backbone movements.

To study PLS, MCCE protonation microstates identify the protonation state of each residue in the protein [56]. MC sampling provides the probability of accepted proton distributions and how these changes when the input structure changes. This procedure has been used to investigate the PLS of the b-type CcO as well as in the quinone loading site in bacterial photosynthetic reaction centers [55, 56].

Here, we explore the PLS in the E-channel of *T. thermophilus*. The study relies on an earlier identification of clusters of amino acids that are highly interconnected, leading across the protein from N- to P-side [10]. Sparse hydrogen bonds connect the clusters. The hypothesis explored here is that a cluster can form a PLS. MCCE is used to calculate the Boltzmann distribution of protonation states in the E-channel in multiple snapshots from MD trajectories carried out with menaquinone (MQ) and menaquinol (MQH_2_) bound or with the apo-proteins. A cluster on the N-side and another in the protein core are found with different net charges in different snapshots, indicating each can bind and release a proton to serve as a PLS. The microstate analysis allows us to determine which residues bind protons showing there are multiple tautomeric states where the protein has the same net charge but different proton distributions in each PLS. There is a strong anticorrelation of proton binding to the N- and P-side cluster (Fig. 7). The central cluster tends to load protons when quinone is bound while the N-side cluster is more likely to be loaded in snapshots from the apo or MQH_2_ bound trajectories.

## 2. Methods

### 2.1. Preparation of the structure for MCCE

The calculations start with the *Thermus thermophilus* apo-Complex I crystal structure (PDB ID: 4HEA) at 3.3 Å resolution [51]. MQ and MQH_2_ bound structures were generated by docking the decyl ubiquinone from the structure 6I0D into the 4HEA structure, modifying the head group and then relaxing the loop from Nqo6 [55–70] (see the details SI.1.1). Earlier studies of the E-channel proton transfer pathways [10] and the motions and orientation of the protein [51, 53] analyzed the trajectories used here.

The subunits are identified using the *T. Thermophilus* numbering. Table 2 and Table SI.2 provide the translation to the chain designations in the 4HEA PDB file and Table SI.1 has the nomenclature of the core subunits of Complex I in different organisms.

MD simulation is carried out on the full membrane embedded apo, MQ and MQH_2_ bound Complex I [51]. For MCCE calculations of the E-channel, only the subunits Nqo4, 7, 8, 10 and 11 are included (Fig. 1). All MD water molecules are removed and substituted with a continuum solvent with a dielectric constant of 80 with 150 mM salt and the protein is given a dielectric constant of 4. A 30 Å low-dielectric rectangular region is created to mimic the hydrophobic portion of the membrane using the IPECE program (See the details in method SI.1.2) [58].

Snapshots were selected for MCCE input in several ways. Every 10^th^ snapshot in each trajectory was subjected to MCCE analysis. Following the MCCE calculations, we identified regions within the MD trajectories where the proton binding stoichiometry changes. Snapshots were added to include more examples with different proton loading. Additionally, snapshots from each trajectory were binned using the MDanalysis [59, 60] and TTClust [61] programs, based on varying residue distances and snapshots were added to ensure that each structural class qA represented. MCCE simulations were subsequently performed on 40 snapshots from the MQ, 20 from the MQH_2_ and 20 from the apo-trajectories.

### 2.2. MCCE calculations on E-channel of Complex I

The MCCE calculations use the ‘isosteric’ conformer choices. The backbone position is fixed given the MD snapshot input. MCCE samples the ensemble of states that differ in the protonation state of Asp, Glu, Tyr, Cys, Arg, His and Lys, hydroxyl and neutral His tautomers and Asn, Gln side chain amide orientation [57, 62]. The MCCE calculations are identical to those used for the analysis of the E-channel hydrogen bond network with one exception. Previously explicit water molecules were retained from the MD snapshots as these are essential to complete the proton transfer paths [10]. Here, implicit water is used as this provides a more reproducible calculation of protonation states with less computational effort [58]. SI.1.3 provides additional details about MCCE force field.

### 2.3. Microstate analysis

MCCE makes side chain and ligand conformer choices for position and charge states. A microstate, which is one conformer choice for each group, is subjected to Metropolis-Hasting MC sampling (additional details in method SI.1.4). All accepted microstates are saved for analysis [56]. Protonation microstates define the charge of each acidic and basic residue [56]. A given protonation microstate can exist in numerous conformations. Protein tautomers are protonation microstates with the same total charge but different proton location [56].

### 2.4. Energy difference between states with different net charge

The free energy difference between any two states can be obtained if we know their probability in the Boltzmann ensemble. Here the probability of all tautomers, i.e. microstates with the same charge, are combined. The free energy difference for a cluster of residues in microstates with an extra proton bound (loaded) or unbound (unloaded) is:

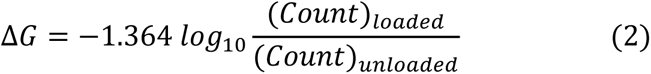

Here, (*Count*)_*loaded*_, is the total count of accepted microstates with an extra proton bound, and (*Count*)_*unloaded*_, is the total count of those with one fewer proton. The unloaded state is the reference here and this calculation is carried out given the microstate distribution for MCCE calculations from a single MD snapshot.

## 3. Results

The goal of this paper is to identify the proton loading site (PLS) of the E-channel in Complex I. Determining whether the PLS proton binding differs in trajectories equilibrated with oxidized or reduced MQ, or in the absence of MQ may help untangle the sequence of proton transfers through the E channel. The PLS are identified by changes in the Boltzmann distribution of ionization states in clusters of residues along the proton transfer path using multiple MD snapshots through a unique MC sampling method [63]. Although the entire protein is included in the MD trajectory, the analysis focused specifically on subunits Nqo4, Nqo7, Nqo8, Nqo10, and Nqo11, which are essential for complex proton transportation and form the unique E-channel (Fig. 1b).

### 3.1. Microstate Energy Distribution

#### 3.1.1. Many Protonation Microstates are Found in the E-channel

The five subunits making up the Complex I E-channel analyzed here contain 1,135 residues with 224 protonatable residues (Asp, Glu, Tyr, His, Lys, Arg). In an MCCE calculations of a representative MQ-docked snapshot at pH 7 there are >600,0000 unique conformation/protonation microstates accepted. In solution, at pH 7 Asp, Glu, the C-terminus, Lys and Arg would be ionized with neutral Tyr, His and the N-terminus in a mixture of charge states. Here, 74 of the residues are found to not be in their ‘expected’ charge state in accepted microstates and many are in protonation states in different microstates. There are >25,000 unique accepted protonation microstates, with ∼240 microstates making up 50% of the total ensemble and ∼8,000 needed to contain 90% of the total (Table 1 and Table SI.3). Thus, MCCE finds every snapshot in many possible protonation states.

**Table 1.** Residues and microstates in E-channel.

| Count | Mix loaded-unloaded frames |  |  |
| --- | --- | --- | --- |
|  | E-Channel | Central Cluster | N-side cluster |
| Asp, Glu, His, Tyr, Arg, Lys | 224 | 15 | 30 |
| Residues changing charge | 74 | 6 | 5 |
| Average ensemble charge | -3.2±0.5 | -1.5±0.1 | -3.4±0.1 |

| Number protonation microstate count |  |  |  |
| --- | --- | --- | --- |
| All | 25,402±970 | 36±10 | 25±5 |
| 90% of ensemble | 8,200±126 | 7±2 | 3±0 |
| 50% of ensemble | 247±56 | 2±0 | 1±0 |

**Table 2.** Residues loading and unloading protons in the Central and N-side clusters in Nqo4, 7, and 8.

| Cluster name | Active residues | *T. Thermophilus* Subunits<br>(4HEA chain ID) |
| --- | --- | --- |
| Central cluster | D72, E74 | Nqo7(P) |
|  | E163, E130, E213, H251 | Nqo8(Q) |
| N-side cluster | H38, E50, **<u>E216</u>**, **<u>E51</u>**, **<u>H58</u>** | Nqo4(E) |
|  | H60 | Nqo7(P) |
|  | E225 | Nqo8(Q) |

MCCE calculated microstates in the complete E-channel (subunits Nqo4, 7, 8, 10, and 11), and in the Central and N-side clusters at pH 7. Data from 10 representative MQ-docked snapshots with the Central Cluster average Central Cluster charge of −1.49 and thus in a mixture of loaded and unloaded microstates shown here. See Table SI.3 for results from different snapshots. Residues in the Central and N-side Clusters were identified in the previous analysis of the proton transfer paths [10]. The active residues in the Central and N-side Clusters are listed in Tables 2 and all cluster residues in Table SI.2.

Figure 3a shows the distribution of >25,000 protonation microstates in one frame from the MQ-trajectory (Table 1). Each dot is a unique protonation microstate. The dots in a column are tautomers, sharing the same charge but with different residues protonated. Some protonation microstates are found in >100,000 accepted microstates while others are accepted <10 times. In a microstate, each protonatable group has a defined charge of 0 or ±1, so microstates have integer charges. Accepted microstates in the E-channel have charges ranging from −9 to +3 with the most probable charges between −6 and 0, centered between −2 to −4 (Fig. 3a). The Boltzmann averaged ensemble is formed of microstates with different charges so has a non-integer value. The ensemble average charge for this representative snapshot is −3.2. See Fig. SI.1 for the distribution of microstates in apo and MQH_2_-docked snapshoots. The detailed explanation of the different clusters, their residues, charge states, and protonation microstates are given in the following subsections.

### 3.2. Central Cluster Loaded and Unloaded States

#### 3.2.1. Microstate Analysis of the Central Cluster

With so many residues in multiple protonation states there is a combinatoric explosion in the number of microstates. However, this can be tamed by looking at individual, connected regions of the protein. Previous mapping of the proton transfer paths through the E-channel identified six clusters of highly interconnected residues (Fig. 2) [10]. Analysis of individual MD snapshots identified residues in clusters, where each is connected to at least 6 other residues either directly or through four or fewer water molecules in different MCCE conformation/protonation microstates. Clusters are then connected by a few hydrogen bonds. The clusters form natural regions to be analyzed independently. The hypothesis is that the protonation states will come to equilibrium in each highly interconnected cluster. Proton transfer between clusters can be gated by the limited inter-cluster connections.

**Fig. 2.**
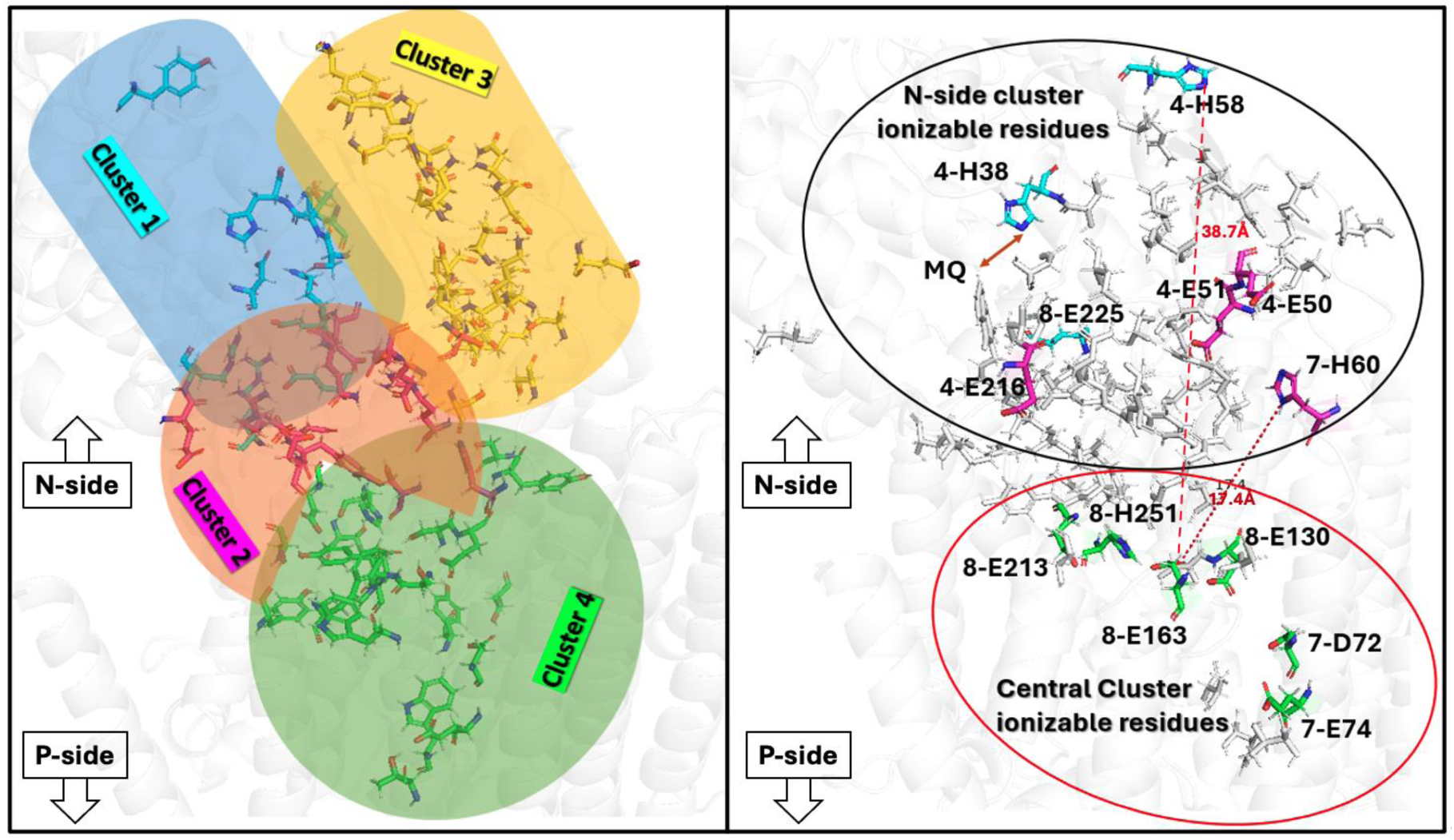
(a) All the cluster residues from a comprehensive analysis of H-bond network by Khaniya et. al. [10]. Different colors show the cluster regions (cluster numbers 1-6 were assigned there): N-side Cluster: (1: cyan; 2: magenta, 3: yellow) and Central Cluster (4: green). Not shown: P-side cluster (cluster 6) below the Central Cluster and Cluster 5: between Cluster 3 and 4. (b) Residues that are active in PLS proton binding and release in: Top Circle: **N-side cluster**; Bottom Circle: Central Cluster. Residue labels: *T. thermophilus* subunit number designation, one letter amino acid code, residue number. See Table SI.1 for glossary of chain designation in Complex I from different species.

On the N-side there is a cluster surrounding the quinone binding site (cluster 1 [10]) with 11 residues (Fig. 2a). This is flanked by two clusters connected to the surface (clusters 2 and 3) with 37 residues (Fig. 2a) [10]. These three clusters will be grouped together as the N-side cluster. The N-side cluster is linked to the 26 residues in the deeply buried Central Cluster (cluster 4) (Fig. 2a). A smaller, isolated cluster 5 was identified. The P-side exit cluster (6) has 32 residues, including several exposed on the P-side surface. The proton path was found to extend from the quinone binding site through the Central Cluster, with rare connections to the P-side exit cluster.

#### 3.2.2. The Central Cluster Net Charge

The Central Cluster has seven acidic residues (Asp, Glu), three bases (Lys, Arg, His), thirteen Grotthuss competent residues (Ser, Thr, Tyr) and three Non-Grotthuss polar residues (Asn, Gln, Trp) (Table 2 and Table SI.2). All these residues are buried. MCCE is carried out on multiple snapshots extracting the five E-channel subunits. First the Boltzmann average net charge is determined for the Central Cluster in each snapshot. The Central Cluster charge in snapshots from the MQ docked trajectory ranges from −2.08 to −1.00 (Fig. SI.2a). Thus, different snapshot conformations have different proton affinities. Snapshots with an average charge from −2.08 to − 1.80 are considered unloaded; −1.29 to −1.00 are loaded and those with a net charge from −1.79 to −1.30 are mixed loaded-unloaded (Fig. SI.2a). The non-integer cluster net charge reflects the distribution of individual microstates with different charges. Those with a charge near −2 have mostly unloaded microstates (Fig. 2d); those with a net charge near −1 mostly have microstates with a proton bound (Fig. 2b); and the snapshot with a cluster charge of −1.49 has a nearly equal mixture of loaded and unloaded microstates in the ensemble (Fig. 2c).

Active residues change protonation in different microstates. One letter amino acid code plus residue ID. The underlined residues in the N-side cluster are surface exposed and bold residues are in the quinone containing, peripheral domain. All Central and N-side cluster residues from reference [10], including those that have the same charge in all accepted microstates, are reported in the Table SI.2

Each trajectory has a different proportion of snapshots with different degrees of proton loading. The Central Cluster is almost always unloaded in snapshots from trajectories with an empty quinone site or with MQH_2_-bound. A few snapshots have mixed loaded and unloaded microstates, but no snapshot has mostly loaded microstates. In contrast, the trajectory with MQ bound has similar number of snapshots that are mostly loaded, mostly unloaded, or with a mixture of loaded and unloaded microstates (Fig. SI.2a).

The energy difference between the proteins in different charge states can be determined given the relative probability of microstates of different charges (Equation 2). We can see here the system is not very deeply trapped with or without a proton bound. The free energy difference between states with –1 or −2 net charge is only 1.4 kcal/mol for the snapshots defined as being loaded and 1.9 kcal/mol for those designated unloaded (Table 3).

**Table 3.** Average free energy of proton binding in Central Cluster (Kcal/mol).

| Frame Types | Avg. Charge | Microstate charge |  |  |  |  | Number snapshots |
| --- | --- | --- | --- | --- | --- | --- | --- |
|  |  | 0 | -1 | - 2 | -3 | -4 |  |
| Loaded | -1.10 | 3.2±0.9 | **<u>-1.4±0.5</u>** | 0 | 3.8±0.8 | >7 | 12 |
| Mixed | -1.49 | 5.0±0.8 | 0.3±0.2 | <u>0</u> | 3.1±0.9 | 5.6±1.5 | 12 |
| Unloaded | -1.99 | 5.5±0.9 | 1.9±0.3 | 0 | 2.5±0.9 | 6.3±1.2 | 9 |

Microstate probability obtained from loaded (average charge −2.08 to −1.80), mixed (−1.79 to - 1.30) and unloaded (−1.29 to −1.00) snapshots. The state with a −2 charge is the reference state. The energies are calculated given the population of microstates with each charge in the ensemble using equation 2. The lowest energy charge state is underlined.

#### 3.2.3. Central Cluster PLS Protonation Microstates: Tracking the Proton Movement

The unique Central Cluster residue protonation microstates were found. Here all conformations and protonation microstates of other residues are counted in one group as long as the Central Cluster residue protonation states are the same. By restricting the analysis to the 15 Central Cluster protonatable amino acids we avoid the combinatoric explosion (Figure 2b, c, d). Now there are 30-40 unique protonation microstates for the Central Cluster. Only 4-5 microstates make up 90% of the ensemble (Table 1 and Table SI.3).

In the Central Cluster Lys and Arg are always ionized, and Tyr is always neutral. Thus, only six Central Cluster residues 8-E163, E130, E213, and H251, and 7-D72, and E74 have different protonation states in the accepted microstates (prefix is the Nqo subunit number) (Table 2). Microstates are found with a charge ranging from −4 to 0. Those with the charges −1 and −2 are the most probable and are considered to be loaded and unloaded (Fig. 3b, c, and d).

**Fig. 3.**
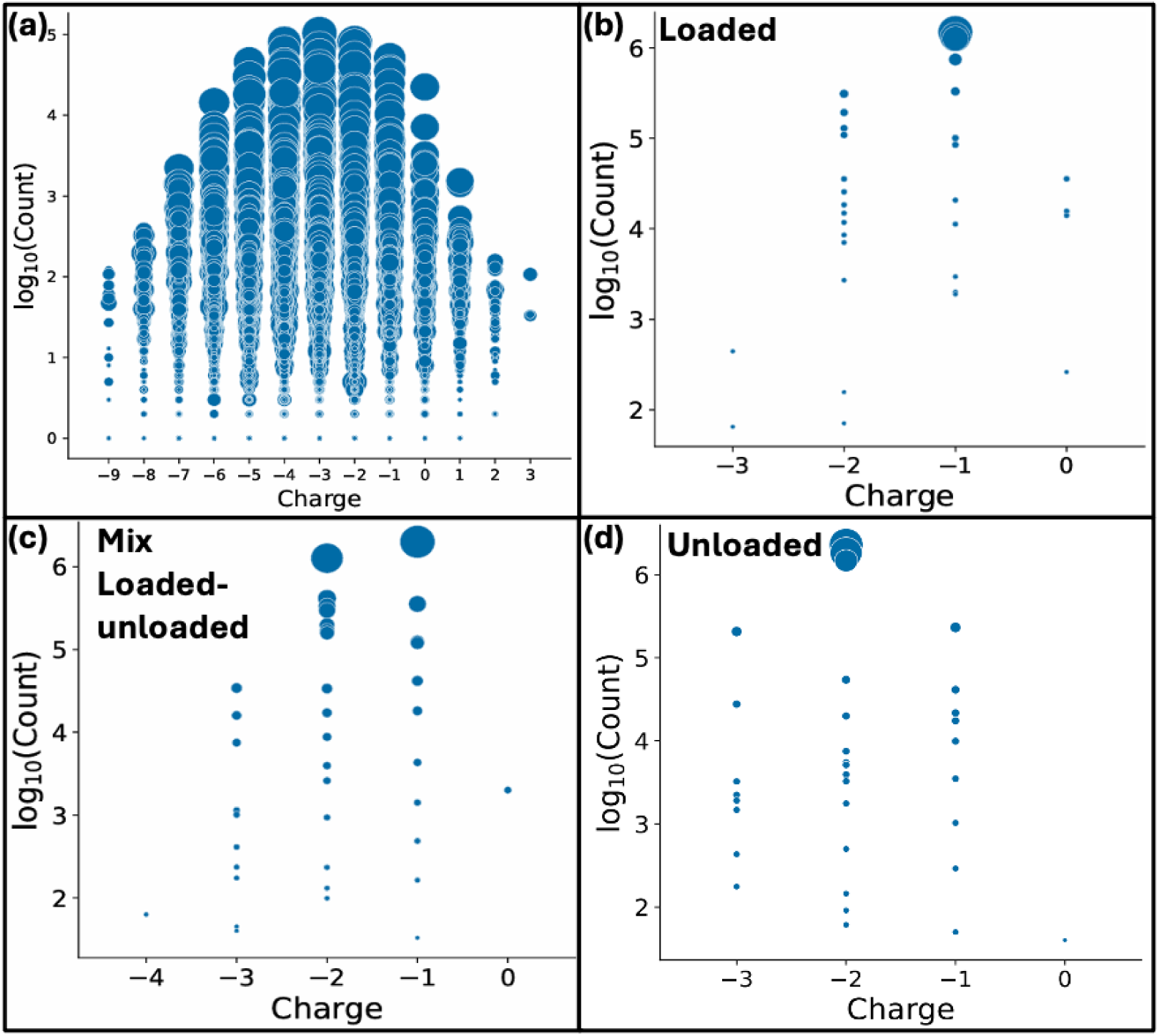
Distribution of unique protonation microstates for three snapshots chosen by the summed charge of the Central Cluster residues. Each dot gives the probability as log_10_(count) and the summed charge of one protonation microstate. Dots in a column are protein tautomers. Dot size indicates the range of energies of the conformational microstates that are found in this protonation state. (a) Five E-channel subunits, average ensemble charge of −3.2. (b-d) Distribution of Central Cluster microstates in snapshots with average net charge of (b) −1.10 (loaded), (c) −1.49 (mixed loaded-unloaded), and (d) −1.99 (unloaded). These microstates differ in the protonation of the six Central Cluster residues that have different charges in different accepted microstates.

We will describe here the results from three representative snapshots, one mostly unloaded (charge −1.99), mostly loaded (−1.11) and one mixed loaded-unloaded (−1.49) (Fig. 3b, c, and d). The microstates described here predominate in 85% of snapshots (Fig. 4c), however their exact probabilities vary. Some snapshots do have other microstates at high probability, which are shown in Table SI.4. The ensemble for each snapshot includes microstates with different net charges, as expected given the modest free energy difference between loaded and unloaded states (Table 3). For example, in the loaded frame the three highest probability microstates have a net charge of −1. Microstates with a charge of −2 and even −3 and 0 are found with low probability (Fig. 3b). In the unloaded frame, microstates with a −2 charge represent 90% of the total (Fi. 3c). The mixed loaded-unloaded frame has an ≈50:50 loaded-unloaded protonation microstates with a −1 and −2 charge (Fig. 3d).

**Fig. 4.**
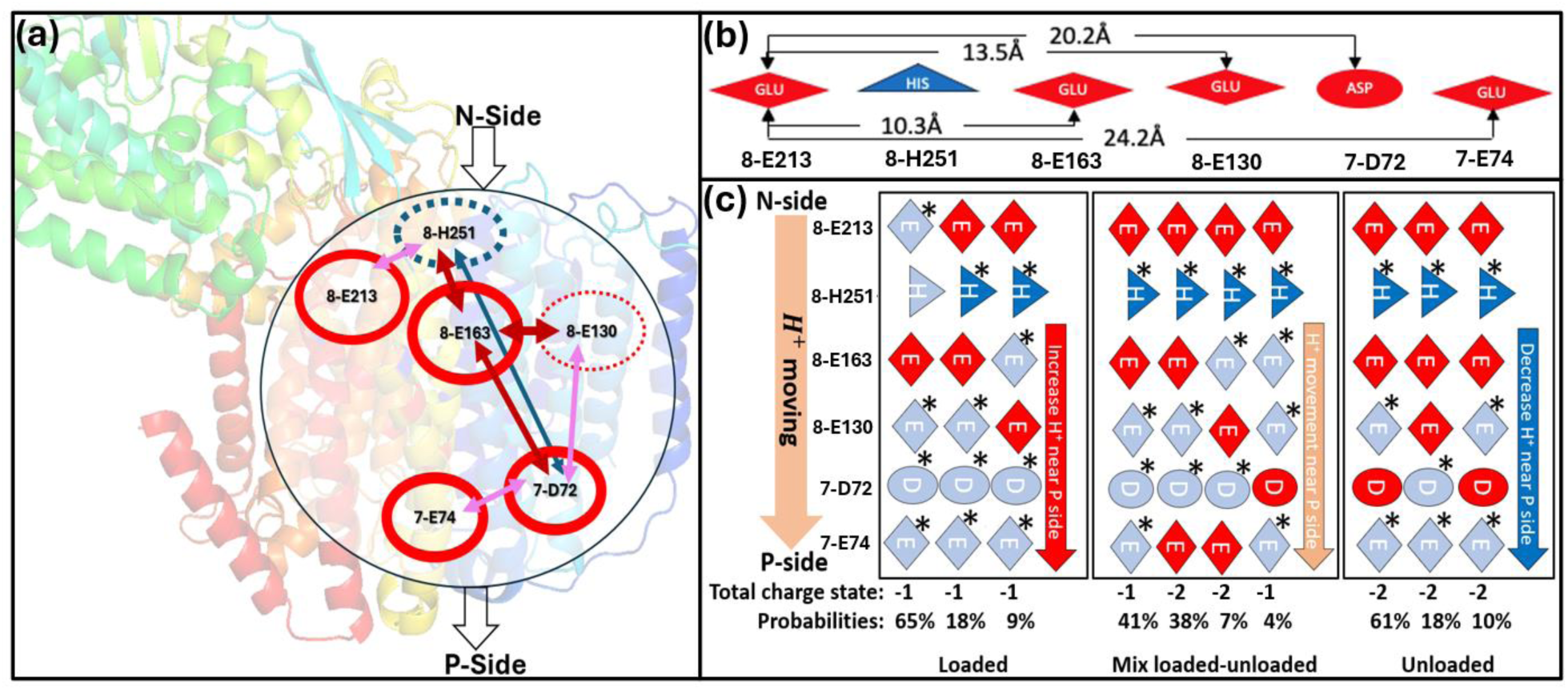
Protonation of the Central Cluster active residues in microstates from loaded, mixed and unloaded snapshots. (a) The Central Cluster residues whose protonation states change between the high probability loaded and unloaded microstates. The arrows indicate the coupling between the residues (Fig. 5). Red arrow: Negatively correlated protonation changes (when one residue is more ionized the other is less); Blue arrow: positive correlation. Arrow thickness indicates correlation strength. Circles around residue number reflect residue conservation in a multiple sequence alignment (Fig. 8). Solid: highly conserved; dashed: lower conservation. Red circles: acids; Blue: bases. (b) Distances between the residues. (c) Highest probable protonation microstates found in MCCE analysis of a snapshot where the Central Cluster is loaded (left), mixed loaded-unloaded (middle) and unloaded (right). The microstates are shown vertically from the N- to the P-side (shapes: diamond: Glu, oval: Asp, triangle: His) Dark red: Glu^-^ or Asp^-^; dark blue: His^+^; Gray: neutral. * Protonated residue.

Figure 4 describes the predominant Central Cluster microstates in three snapshots. The Central Cluster extends over 24Å from 8-E213 near the N-side to 7-E74 nearer the P-side (Fig. 4b). A schematic indicates the location of the proton in each microstate with an asterisk identifying sites with a proton. The net microstate charge as well as its probability in the full ensemble are provided. In unloaded frames the P-side residues 7-D72 and 7-E74, which are within 5.0 Å of each other, are mostly deprotonated (Fig. 4c). In loaded states a proton is almost always bound by either or both acids (Fig. 4). Thus, the proton moves towards the P-side of the loaded Central Cluster (Fig. 4). In contrast, the mixed loaded-unloaded snapshots tend to bind protons nearer the N-side on 8-E163 and 8-E130 (Fig. 4). Thus, the loaded Central Cluster stabilizes protons nearer the P-side of the extended cluster, poising them for release.

### 3.3. Correlation of Central Cluster Residue Protonation

One significant advantage of having access to all microstates lies in the ability to uncover correlations amongst residues. The Pearson weighted correlation of protonation states of the Central Cluster residues was determined (SI. Equation 1) [56]. A positive correlation suggests that the two groups are more likely to be ionized simultaneously, whereas a negative correlation indicates that the ionization of one group decreases the likelihood of the other group being ionized (see SI.1.5). All the active groups in the Central Cluster influence each other. Residue 8-E163 is pivotal, as its ionization is negatively correlated with 7-D72 and 8-E130 and positively correlated with 8-H251 (Fig. 5). Likewise, 8-H251 is positively correlated with 7-D72 and 7-E74. Thus, a proton tends to be on the His on the N-side or 24 Å away on one of the P-side acids. This highlights the long-range coupling within this cluster. The correlation coefficient is never close to one, indicating that these correlations are preferences rather than rules. Correlation analysis for loaded and unloaded snapshots can be found in Figure SI.2.

**Fig. 5.**
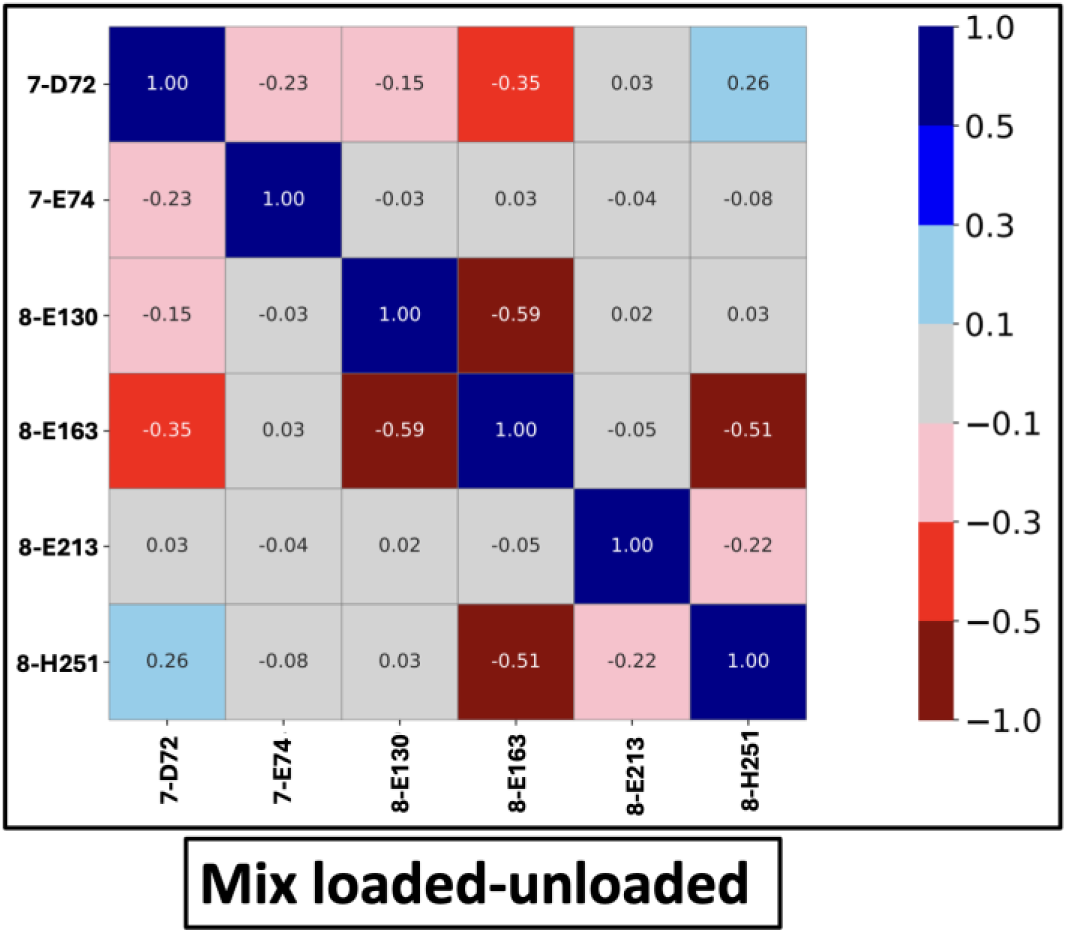
Pearson weighted correlation coefficient (SI. Equation 2) for Central Cluster residue protonation in a mixed loaded-unloaded snapshot. Only residues with an absolute correlation value ≥0.1 with at least one other residue are included. Each square gives the correlation strength between two residues. The color denotes correlation values: dark red (−1.0 to −0.5), red (−0.5 to −0.3), pink (−0.3 to −0.1), light gray (−0.1 to 0.1), sky blue (0.1 to 0.3), blue (0.3 to 0.5), and dark blue (0.5 to 1.0). The correlations shown here are the source of the weight and color of the arrows in Fig. 4a.

The two adjacent P-side acids, 7-D72 and 7-E74 are 5 Å apart from each other. 7-D72 has been previously discussed as part of a PLS [9, 19, 28, 54, 64]. Here we find that both contribute, and their ionization is anticorrelated (Fig 5 and Fig. SI.3). They are both predominately neutral in loaded structures; however, the probability of 7-D72 and 7-E74 being ionized in higher energy microstates is similar (Fig. 4). In the mixed loaded-unloaded snapshots 7-E74 is more likely to be ionized, whereas in the fully unloaded snapshots 7-D72 is ionized while 7-E74 retains the proton.

### 3.4. The N-side Cluster

Three clusters were identified in the analysis of the H-bond networks close to the N-side of the E-channel, previously designated clusters 1, 2, and 3 [10] (Fig. 2). There is five Asp, nine Glu, one Lys, six Arg and three His in the combined N-side cluster (Table SI.2). All Arg and Lys remain ionized and Tyr neutral. However, 4-H38, 4-E50, 4-E51, 4-E216, 7-H60, and 8-E225 are in different protonation states (Table 2). Among them 4-E58, 4-E51, and 4-E216 are in the peripheral arm while 4-E51, 4-E50, 4-E216, 8-E225, and 7-H60 are near each other, just above the membrane surface (Fig. 2).

Residues 4-H38, D139, and H58 are not included in the calculation of the N-side cluster charge. While these residues are connected to the other residues via the hydrogen bond network, they are far from the membrane and thus less likely to inject protons into the E-channel. MQ is bound to 4-Y87 and 4-H38, while 4-D139 is bound to 4-H38 [54, 65–67]. 4-D139 is always ionized and 4-Y87 neutral. 4-H58 is ≈40 Å further away from the membrane in the periplasmic arm (Fig. 2b). 4-H38 and 4-H58 do change protonation in our analysis, but their ionization is not correlated with each other. The correlation of 4-H38 and the other N-side cluster residues are discussed below.

#### 3.4.1. The N-side Cluster Net Charge

The average N-side cluster charge varies between −3.8 and −2.6 in different snapshots, identifying it as a second PLS. A loaded cluster has a charge of −3 while it is −4 in the unloaded state (Fig. SI.2b). The large negative charge is supported by the positive electrostatic potential from the remaining protein. The N-side cluster charge in MQ trajectory snapshots ranges from - 3.8 to −2.9 indicating some snapshots are loaded and others unloaded (Fig. SI.2b). The MQH_2_ and Apo trajectory snapshots are mostly loaded with the total charge between −3.1 and −2.7 in the apo and −2.9 and −2.6 in the MQH_2_ trajectory. This implies the proton-donating residues in the Complex I MQ sites get protonated while MQH_2_ still remains bound to the cavity. The distribution of snapshots with different average charges is shown in (Fig. SI.2b).

#### 3.4.2. N-side Cluster PLS Protonation Microstates: Tracking the Proton Interacting Residues

There are five N-side cluster residues whose protonation states differ in highly probable microstates. In the highest probability unloaded microstate, which makes up 61% of the represented unloaded snapshot shown in (Fig. 6b) none of the residues has a proton. In the most probable loaded microstates 7-H60, the residue closest to the Central Cluster, is protonated (Fig. 6a). Lower probability loaded microstates have the proton on either 4-E216, 4-E50 or 4-E51. There is less correlation amongst the protonation states of the N-side residues than what was found for those in the buried Central Cluster PLS. The strongest correlation is between 4-E50 near the N-side surface and 7-H60, which is the residue closest to the Central Cluster (Fig. SI.4).

**Fig. 6.**
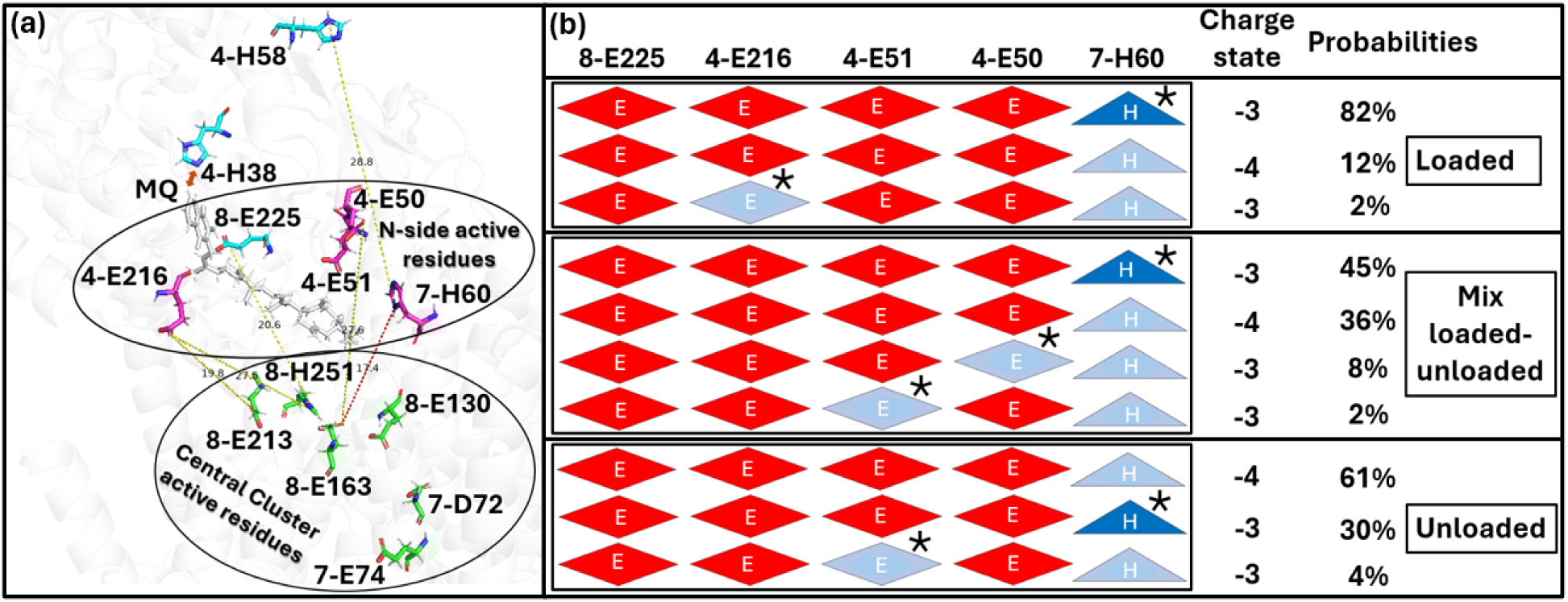
N-side cluster protonation microstates. (a) N-side and Central Cluster active residues. (b) Most probable protonation microstates found in representative snapshot with N-side cluster loaded (top), mixed loaded-unloaded (middle) and unloaded (bottom). The microstates are shown horizontally with the same residue shape and color designation found in Fig. 4.

**Fig. 7.**
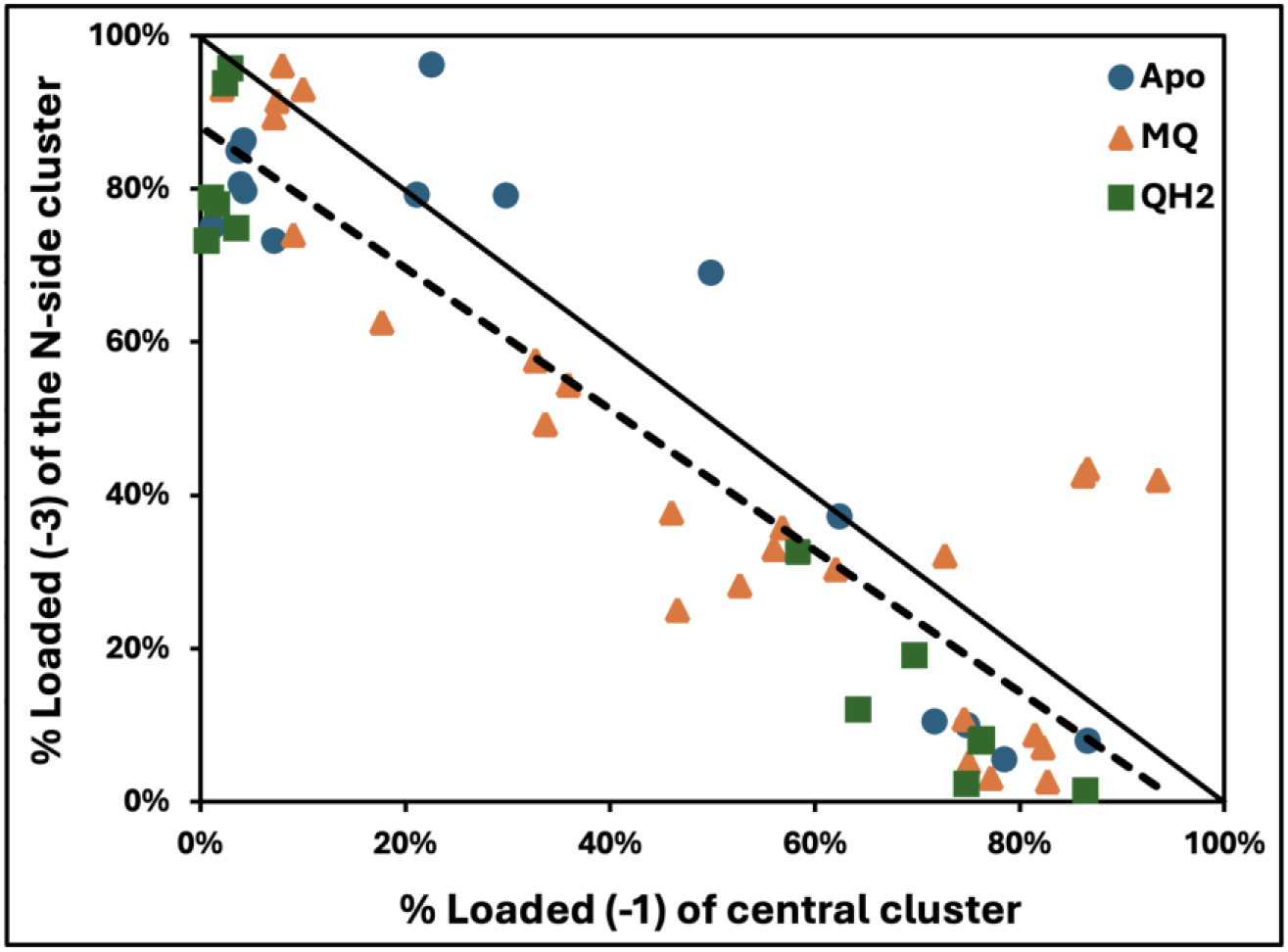
Correlation of Central and N-side PLS loading. A dot shows the percentage of loaded Central Cluster microstates (charge state: −1) vs. N-side cluster (charge state: −3) in individual snapshots from the Apo (blue), MQ (orange), and MQH_2_ (green) docked MD trajectories. Dashed line: Best fit **slope −0.92 and R² 0.82; Solid line: y = -x.**

#### 3.4.3. Correlation Between Central and N-side Cluster Loading

The average charge of the Central and N-side clusters was compared for each individual snapshot. Protonation of the two clusters is strongly anti-correlated (Fig. 7). In snapshots from the MQH_2_ trajectory the N-side cluster tends to be loaded and Central Cluster unloaded. Snapshots from the Apo and MQ trajectories are more diverse showing a range of loading for the two PLS, with the apo snapshots favoring a more loaded N-side cluster. On average the two PLS together hold on average ∼0.9 protons. There are however a small number of snapshots where there are from 1.2-1.4 protons loaded between the N-side and Central Clusters.

### 3.5. Other Buried Clusters

#### 3.5.1. Second Buried Cluster

A small cluster, designated cluster 5, was identified in the study of the proton transfer paths. There were no connections between this cluster and any other cluster even when additional water molecules were added to aid connectivity [10]. The three ionizable residues in the cluster are 10-E32, 8-Y43 and 8-Y59. 10-E32 changes charge in the different snapshots, especially in snapshots from MQ and apo trajectories. It is mostly neutral in those from the MQH_2_ trajectory. However, its ionization is not correlated with the Central Cluster charge (Fig. SI.5).

#### 3.5.2. The P-side Cluster

The Central Cluster is 25 Å from the P-surface. A P-side cluster, which is largely parallel to the membrane surface, was identified. Only very rare connection were found between the Central and P-side cluster in the study of the proton transfer paths [10]. The total charge of the cluster is - 1 and rarely changes in our simulation, even when we look at low probability microstates. The constant charge of the cluster indicates the alternative protonation microstates are >6 kcal/mol higher in energy. However, there are fluctuations in the ionization of 10-E5 or 8-K190 in several apo state snapshots. These changes either lead to low probability cluster binding or releasing of a proton.

### 3.6. Interaction of 4-H38 and 4-D139 in MQ Binding Region

The residue 4-H38 is a ligand to the MQ and can make a hydrogen bond to 4-D139. A break of the His-Asp hydrogen bond on quinone reduction has been proposed as a trigger for proton pumping [54]. The parent structure, 4HEA, has them close together. In the apo trajectory, the pair maintain a short distance near ∼2.6Å until the last frames where they begin to move apart (Fig. SI.6a, and c). In the MQH_2_ trajectory the pair quickly moves to be ∼4.7Å apart. In the MQ-trajectory the distance between the His and Asp starts near ∼2.3 Å and fluctuates, expanding beyond 8 Å.

During all MD trajectories, His is fixed neutral and Asp ionized (Fig. SI.6). Despite this the His takes different charge states in the MCCE calculations, while the Asp remains ionized. In the MCCE analysis of MQH_2_ snapshots, where the distance between the pair is intermediate and stable, the His is almost always neutral, perhaps as the nearby MQH_2_ is protonated. In contrast, in the Apo snapshots, the Asp-His distance remains short, but the average His ionization in different snapshots ranges from zero to 60% ionized, as there is no quinone for it to interact with. The distance varies in the MQ trajectory. The His is largely ionized when it is closest to the Asp so that it can hydrogen bonded to both quinone and the ionized Asp; while it is more likely to be neutral when separated from the Asp. However, the correlation of ionization with distance is modest.

## 4. Discussion

Proton pumps require proton transfer paths connecting PLS, which are sites for temporary loading and unloading of protons. Here we examined the proton loading behavior of highly interconnected regions along the previously identified [10] proton transfer path through the E-channel in Complex I. The assumption underlying this study is that proton distribution within each cluster will be at equilibrium with the structure. Implicit assumptions are that protons bound to sites off the path will not be transported so they cannot be part of PLS; and that gating can be achieved either by breaking the sparser inter-cluster connections or by making the PLS proton affinity so high or low that protons cannot move through it [21].

### Residues in the PLS

Analysis of the Boltzmann distribution of microstates generated in MC sampling shows competing, high probability protonation microstates. We found two PLS, each with multiple residues stretching over many angstroms. Thus, the PLS are not individual residues but rather each an extended, interconnected group of residues. The proton affinity of the cluster determines PLS loading. The loading of the two PLS are anti-correlated suggesting that a proton bound to the N-cluster may be transferred to the Central Cluster. On average ≈90% of the two coupled PLS have a proton bound (Fig. 7).

There have been earlier studies that pointed to residues of interest in the E-channel identified by inspection of the structure plus consideration of residue conservation [6, 9, 13, 19, 20, 41, 42, 54, 64–66, 68–70]. Some studies calculated changes in the average proton affinity of individual residues using traditional methods that deliver uncorrelated, individual site average protonation [6, 9, 13, 19, 20, 41, 42, 54, 64, 66, 68–70]. MC sampling allows an unbiased examination of all residues, identifying additional residues of interest (Fig. 2b). Comparison of individual protonation microstates shows how the proton can move from N- to P-side of the extended PLS clusters.

We find many previously identified residues are involved in proton loading including 7-D72, 7-E74, 8-E130, 8-E163, 8-E213, 8-H251, 10-Y59, and 11-E32 [6, 9, 13, 19, 20, 41, 42, 54, 64, 66, 68–70]. The MC analysis here identifies additional residues in the N-side cluster just above the membrane including 4-E51, 4-E50, and 7-H60. These residues form water molecule mediated, hydrogen bonds and change protonation in highly probable protonation microstates. The identification of the N-cluster as a PLS may support earlier computational analyses [19, 54] that proposed the protonation states of Nqo7 Asp and Glu residues near the solvent interface might be part of proton-conducting water molecule channels.

### Complex vs Simple Pumping Elements

Proton paths and their PLS are being found in two flavors. There are linear paths such as the D-channel in CcO [21, 71] and the antiporter channels in Complex I [6, 9, 13, 69, 72]. These can be dominated by single isolated residue PLS, such as D132 and E286 at the entry and end of the CcO D-channel [15]. In the antiporters, a linear array of 4-5 protonatable residues form a PLS. Linear proton paths and simple PLS can be found by inspection of the structure. Mutation of a single residue can greatly reduce pumping [37–39, 70, 73–75]. The residues along these paths are often highly conserved.

In contrast, complex proton transfer paths have a network of competing routes. The associated PLS are more often clusters of residues. Proton pumps seem to use both simple and complex paths. For example, while protons come in to CcO via linear paths they exit via a complex network on the P-side [21]. The hypothesis is that there are multiple ways to move a proton through the pump. Thus, multiple mutations that block alternative paths may be needed to block activity. This is seen in the proton input path in bacterial reaction centers [55] and found here [76].

### Mutation and Conservation of PLS Residues

In Complex I the antiporter channels are simple while the E-channel is not. Experiments measured the rates of NADH oxidization or quinone reduction that are coupled to proton pumping in mutants of *E. coli* or *P. denitrificans* Complex I. Mutation of individual residues identified here as contributing to the PLS led to diminished but not fully inhibited rates (Table SI.5) [70, 76–78]. For example, 7-E163 appears as a key element in the Central Cluster PLS, but when it is mutated the rate of pumping coupled NADH oxidation is slowed by 75%. However, multiple mutations are more lethal. For example, individual removal of 7-D72 or E74, inhibit activity but when both are changed to non-ionizable residues the reaction is 98% inhibited [76] (Table SI.5).

### Central Cluster Conservation

The conservation of the residues that form the Central Cluster PLS and the residues within 5.0 Å around them was determined (see SI.1.6). The coupled residues 8-E213, and 8-E163 near the N-side and 7-D72, and 7-E74 near the P-side are highly conserved (Fig. 8; top). This finding can support 8-E163 being a key during the proton transport through the E-channel in different organisms (Fig. 4a). In contrast, 8-E130 is often a Ser and 8-H251 can be a variety of polar residues (Table SI.5).

**Fig. 8.**
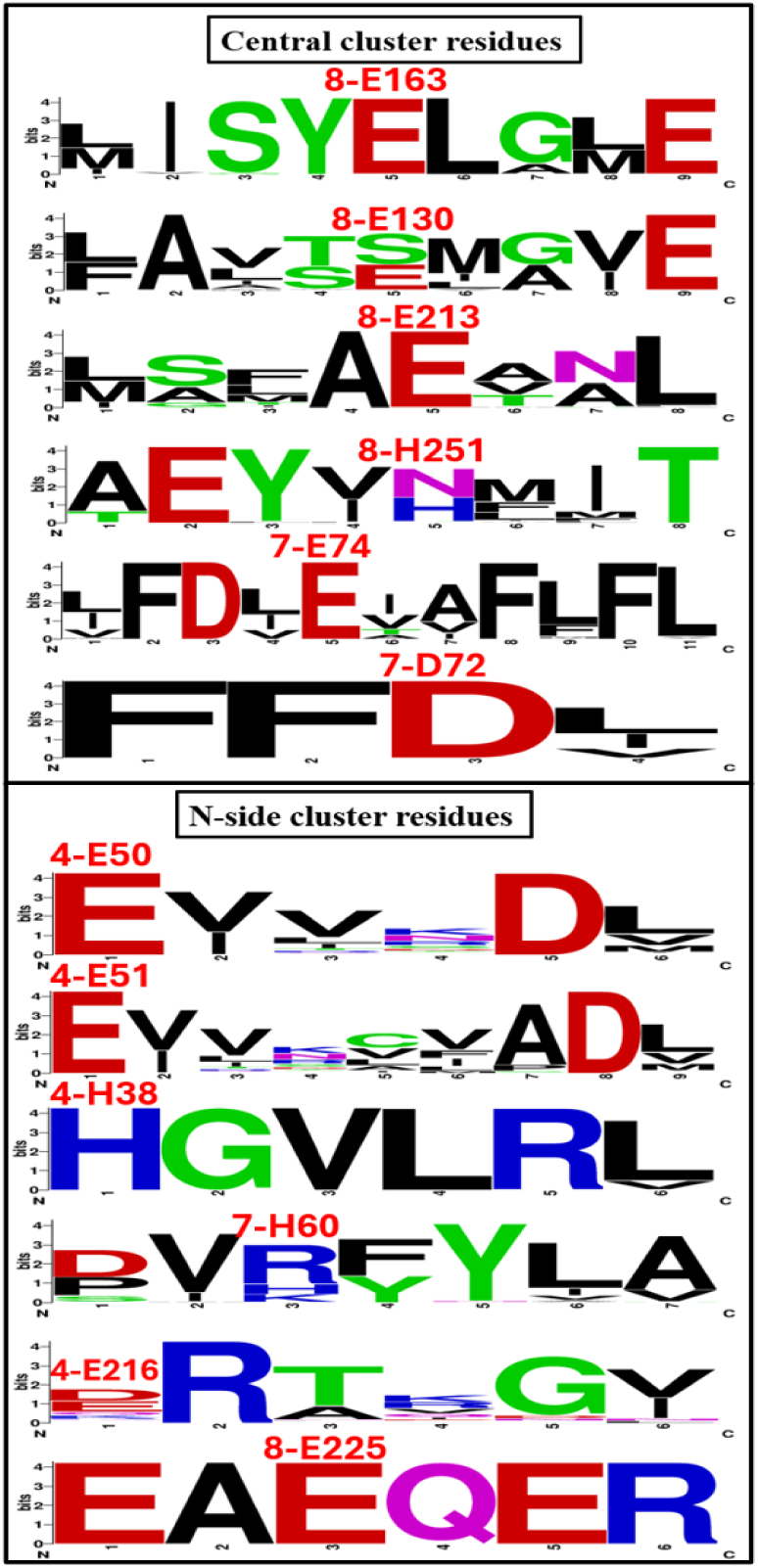
Web-logo [79] representation of multiple sequence alignment of 1000 Complex I sequences for only residues of within 5.0 Å of the active residues in the Central Cluster and the N-side cluster. The surrounding residues in the Central Cluster and the N-side cluster are given in **Table SI.6.** These are in subunits Nqo4, Nqo7, Nqo8, Nq010, and Nqo11 in *T. thermophilus* Complex I. Residues name identified by *T. thermophilus* subunits Nqo4-, Nqo7-, and Nqo8-one-letter residue name, and followed by residues ID shown on top of each key residues on the Web-logo. Their degree of conservation is the source of the circle type (thickness, solid or dashed) in Fig. 4a and Fig. SI.3.

The surrounding residues play two roles in proton transfer. Hydroxyl containing residues are Grotthuss competent; hydrophilic residues can help stabilize bound water molecules needed for proton transfer paths; and the electrostatic potential generated by the surrounding help support the PLS loaded and unloaded charge states. The Central Cluster connects residues separated by over 22.0 Å. The multiple sequence alignment shows polar surrounding residues are often conserved. Overall, the residue types are usually maintained (Fig. 8).

### N-side Cluster Conservation

In the N-side Cluster only 4-H130, 8-E225, 4-E51, 4-E50 and the quinone ligand 4-H38 are highly conserved while 4-H58, 7-H60 and 4-E216 are not (Fig.8 and Table SI.5). Conservation of surrounding, non-protonatable residues is variable, with some well conserved and others not (Fig. 8; top).

### Towards a Unified Model of Pumping

It is challenging to imagine a mechanism that can couple electron transfer in the peripheral arm of Complex I with proton pumping through four channels. Models of pumping consider how the quinone reduction or movement influence the residues between the quinone site and the top of the E-channel, which are in the N-side cluster here [12, 52, 54, 80–82]. Others consider the communication to the antiporter channels, via coupling through the adjacent ionizable residues in the membrane center [9, 12, 82–84]. Models of pumping through the E-channel have been proposed [11, 52], while others suggest no protons are pumped through the E-channel [35].

Here we assume the E-channel pumps a proton. A model of pumping would connect the proton loading and the proton path connections in the different intermediate states. The calculation of PLS loading presented here is carried out on the same trajectories as a previous study of the proton transfer paths (Fig. SI.7) [10]. Thus, we can compare the degree of connectivity and loading of putative E-channel PLS described here.

Snapshots from the MQH_2_ trajectory have the N-side cluster fully loaded, with little proton loading in the Central Cluster and few connections between the two clusters. The quinol would be released and quinone rebound. In the apo trajectory the proton has moved into the Central Cluster and this PLS continues to fill when MQ is bound. The rare connections to the P-side cluster are made in the apo trajectory, indicating this may be the state where protons would be released to the P-side. However, the Central Cluster is most highly filled when MQ is bound, which could also make this a good candidate for proton transfer to the P-side. There are multiple steps going from MQ to MQH_2_ with a first and second reduction and quinone protonation being able to modify the loading of the clusters. Also the quinone and quinol may be transiently bound in different binding sites as it moves from the membrane into the periplasmic arm [80, 81]. However, we do not have information about intermediate structures here.

It should also be noted that the calculations do not show the PLS strongly traps the proton. In all snapshots MCCE finds mixtures of loaded and unloaded microstates with only a few kilocalories separating them. This may be simply due to the limitation of the MD snapshots. Alternatively, it may be a property of Complex I pumping if the strong physical gate found between the Central and P-side clusters ensures the proton only moves when the open gate is triggered.

The P-cluster remains largely inert. The cluster keeps the same protonation state in all snapshots. Thus, microstates with different numbers of protons loaded are at least 6 kcal/mol above those at low energy. Another concern is that there is rarely a path between the Central and P-side cluster. However, pumps are designed not to be opened for transport except when coupled to the redox reactions that power them. Thus, a pathway from the N-side D-channel to the P-side proton exit is not found in any structure or in most MD simulations. A ‘lucky’ MD trajectory found protonation of the P-side PLS triggered the opening of a water molecule-filled cavity in the hydrophobic region near the bridging Glu286, which serves as a single residue PLS. Water entry lowers the Glu proton affinity by more than 5 kcal/mol allowing proton release as well as a connection between the N- and P-sides of the protein [24]. Similarly, in b-type CcO all crystal structures have an extremely high proton affinity for the P-side PLS cluster. MD changed the structure to lower the proton affinity so that the PLS could be characterized [21]. Thus, we posit that the gate opening from the Central to P-side cluster is rare and that our trajectories have a P-side cluster trapped in a single loaded state still missing the gate opened conformation.

## 5. Conclusion

While the antiporter subunit and peripheral arm of Complex I are related to other proteins [6, 9, 11, 44, 46, 47, 69, 85] the E-channel is unique. This channel is challenging to study as there is no linear proton transfer path. MC sampling allows us to identify the many possible paths by mapping the network of hydrogen bond connections [10]. Here MC sampling is used to find and characterize highly interconnected clusters that function as PLS. To identify PLS structures it is required that they differ in their proton affinity. Here MD snapshots in apo, MQ and MQH_2_ states were evaluated. Previously identified clusters were found to change charge. In a cluster at the N-side of the proton, five residues are coupled together to form a PLS. Six residues function in an extended Central Cluster PLS. A small disconnected buried cluster also binds and releases a proton. However, in the structures that are available, the cluster on the P-side of the protein is inert. Many of the residues found to be active in the PLS have been studied previously by site-directed mutation. As found in other complex proton transfer motifs single mutations have modest effects, while double mutations are more lethal. This supports the picture of a complex, robust pumping element where there are multiple alternative ways to travel.

## Acknowledgements

Gunner, Uddin, and Khaniya acknowledge financial support from NSF MCB-2141824. Abhishek Singharoy and Chitrak Gupta acknowledge start-up funds from the SMS and CASD at Arizona State University, and the resources of the OLCF at the Oak Ridge National Laboratory, which is supported by the Office of Science at DOE under Contract No. DE-AC05-00OR22725, made available via the INCITE program.

## References

[1] Mitchell, P., Coupling of phosphorylation to electron and hydrogen transfer by a chemi-osmotic type of mechanism. Nature, 1961. 191(4784): p. 144–148, 10.1038/191144a0.

[2] Abrahams, J.P., A.G. Leslie, R. Lutter, and J.E. Walker, Structure at 2.8 A resolution of F1-ATPase from bovine heart mitochondria. Nature, 1994. 370(6491): p. 621-628, 10.1038/370621a0.

[3] Watt, I.N., M.G. Montgomery, M.J. Runswick, A.G. Leslie, and J.E. Walker, Bioenergetic cost of making an adenosine triphosphate molecule in animal mitochondria. Proceedings of the National Academy of Sciences, 2010. 107(39): p. 16823–16827, 10.1073/pnas.1011099107.

[4] Iwata, S., J.W. Lee, K. Okada, J.K. Lee, M. Iwata, B. Rasmussen, T.A. Link, S. Ramaswamy, and B.K. Jap, Complete structure of the 11-subunit bovine mitochondrial cytochrome bc1 complex. Science, 1998. 281(5373): p. 64-71, 10.1126/science.281.5373.64.

[5] Raven, P.H., Origin of Eukaryotic Cells. Evolution, 1971. 25(4): p. 737–737, 10.1111/j.1558-5646.1971.tb01930.x.

[6] Baradaran, R., J.M. Berrisford, G.S. Minhas, and L.A. Sazanov, Crystal structure of the entire respiratory complex I. Nature, 2013. 494(7438): p. 443-448, 10.1038/nature11871.

[7] Efremov, R.G., R. Baradaran, and L.A. Sazanov, The architecture of respiratory complex I. Nature, 2010. 465(7297): p. 441-445, 10.1038/nature09066.

[8] Ripple, M.O., N. Kim, and R. Springett, Mammalian complex I pumps 4 protons per 2 electrons at high and physiological proton motive force in living cells. Journal of Biological Chemistry, 2013. 288(8): p. 5374–5380, 10.1074/jbc.M112.438945.

[9] Sazanov, L.A., A giant molecular proton pump: structure and mechanism of respiratory complex I. Nature reviews Molecular cell biology, 2015. 16(6): p. 375–388, 10.1038/nrm3997.

[10] Khaniya, U., C. Gupta, X. Cai, J. Mao, D. Kaur, Y. Zhang, A. Singharoy, and M. Gunner, Hydrogen bond network analysis reveals the pathway for the proton transfer in the E-channel of T. thermophilus Complex I. Biochimica et Biophysica Acta (BBA)- Bioenergetics, 2020. 1861(10): p. 148240, 10.1016/j.bbabio.2020.148240.

[11] Berrisford, J.M., R. Baradaran, and L.A. Sazanov, Structure of bacterial respiratory complex I. Biochimica et Biophysica Acta (BBA)-Bioenergetics, 2016. 1857(7): p. 892–901, 10.1016/j.bbabio.2016.01.012.

[12] Kampjut, D. and L.A. Sazanov, Structure of respiratory complex I–An emerging blueprint for the mechanism. Current opinion in structural biology, 2022. 74: p. 102350, 10.1016/j.sbi.2022.102350.

[13] Kaila, V.R., Long-range proton-coupled electron transfer in biological energy conversion: towards mechanistic understanding of respiratory complex I. Journal of The Royal Society Interface, 2018. 15(141): p. 20170916, 10.1098/rsif.2017.0916.

[14] Balashov, S.P., Protonation reactions and their coupling in bacteriorhodopsin. Biochimica et Biophysica Acta (BBA)-Bioenergetics, 2000. 1460(1): p. 75–94, 10.1016/S0005-2728(00)00131-6.

[15] Kaila, V.R., M.I. Verkhovsky, and M. Wikström, Proton-coupled electron transfer in cytochrome oxidase. Chemical reviews, 2010. 110(12): p. 7062–7081, 10.1021/cr1002003.

[16] Wikström, M., Cytochrome c oxidase: 25 years of the elusive proton pump. Biochimica et Biophysica Acta (BBA)-Bioenergetics, 2004. 1655: p. 241–247, 10.1016/j.bbabio.2003.07.013.

[17] Gunner, M., M. Amin, X. Zhu, and J. Lu, Molecular mechanisms for generating transmembrane proton gradients. Biochimica et Biophysica Acta (BBA)-Bioenergetics, 2013. 1827(8-9): p. 892–913, 10.1016/j.bbabio.2013.03.001.

[18] Gunner, M. and R. Koder, The design features cells use to build their transmembrane proton gradient. Physical Biology, 2017. 14(1): p. 013001, 10.1088/1478-3975/14/1/013001.

[19] Kaila, V.R., M. Wikström, and G. Hummer, Electrostatics, hydration, and proton transfer dynamics in the membrane domain of respiratory complex I. Proceedings of the National Academy of Sciences, 2014. 111(19): p. 6988–6993, 10.1073/pnas.1319156111.

[20] Zickermann, V., C. Wirth, H. Nasiri, K. Siegmund, H. Schwalbe, C. Hunte, and U. Brandt, Mechanistic insight from the crystal structure of mitochondrial complex I. Science, 2015. 347(6217): p. 44-49, 10.1126/science.1259859.

[21] Cai, X., C.Y. Son, J. Mao, D. Kaur, Y. Zhang, U. Khaniya, Q. Cui, and M. Gunner, Identifying the proton loading site cluster in the ba3 cytochrome c oxidase that loads and traps protons. Biochimica et Biophysica Acta (BBA)-Bioenergetics, 2020. 1861(10): p. 148239, 10.1016/j.bbabio.2020.148239.

[22] Kaur, D., X. Cai, U. Khaniya, Y. Zhang, J. Mao, M. Mandal, and M.R. Gunner, Tracing the pathways of waters and protons in photosystem II and cytochrome c oxidase. Inorganics, 2019. 7(2): p. 14, 10.3390/inorganics7020014.

[23] Yang, L., Å.A. Skjevik, W.-G.H. Du, L. Noodleman, R.C. Walker, and A.W. Götz, Water exit pathways and proton pumping mechanism in B-type cytochrome c oxidase from molecular dynamics simulations. Biochimica et Biophysica Acta (BBA)- Bioenergetics, 2016. 1857(9): p. 1594–1606, 10.1016/j.bbabio.2016.06.005.

[24] Goyal, P., J. Lu, S. Yang, M. Gunner, and Q. Cui, Changing hydration level in an internal cavity modulates the proton affinity of a key glutamate in cytochrome c oxidase. Proceedings of the National Academy of Sciences, 2013. 110(47): p. 18886–18891, 10.1073/pnas.1313908110.

[25] Wang, P., N. Dhananjayan, M.A. Hagras, and A.A. Stuchebrukhov, Respiratory complex I: Bottleneck at the entrance of quinone site requires conformational change for its opening. Biochimica et Biophysica Acta (BBA)-Bioenergetics, 2021. 1862(1): p. 148326, 10.1016/j.bbabio.2020.148326.

[26] Chung, I., D.N. Grba, J.J. Wright, and J. Hirst, Making the leap from structure to mechanism: are the open states of mammalian complex I identified by cryoEM resting states or catalytic intermediates? Current opinion in structural biology, 2022. 77: p. 102447, 10.1016/j.sbi.2022.102447.

[27] Hirst, J., Mitochondrial complex I. Annual review of biochemistry, 2013. 82: p. 551–575, 10.1146/annurev-biochem-070511-103700.

[28] Wirth, C., U. Brandt, C. Hunte, and V. Zickermann, Structure and function of mitochondrial complex I. Biochimica et Biophysica Acta (BBA)-Bioenergetics, 2016. 1857(7): p. 902–914, 10.1016/j.bbabio.2016.02.013.

[29] Agip, A.-N.A., J.N. Blaza, J.G. Fedor, and J. Hirst, Mammalian respiratory complex I through the lens of cryo-EM. Annual review of biophysics, 2019. 48(1): p. 165–184, 10.1146/annurev-biophys-052118-115704.

[30] Agip, A.-N.A., J.N. Blaza, H.R. Bridges, C. Viscomi, S. Rawson, S.P. Muench, and J. Hirst, Cryo-EM structures of complex I from mouse heart mitochondria in two biochemically defined states. Nature structural & molecular biology, 2018. 25(7): p. 548–556, 10.1038/s41594-018-0073-1.

[31] Djurabekova, A., J. Lasham, O. Zdorevskyi, V. Zickermann, and V. Sharma, Long-range electron proton coupling in respiratory complex I—insights from molecular simulations of the quinone chamber and antiporter-like subunits. Biochemical Journal, 2024. 481(7): p. 499–514, 10.1042/BCJ20240009.

[32] Yoga, E.G., H. Angerer, K. Parey, and V. Zickermann, Respiratory complex I– mechanistic insights and advances in structure determination. Biochimica et Biophysica Acta (BBA)-Bioenergetics, 2020. 1861(3): p. 148153, 10.1016/j.bbabio.2020.148153.

[33] Bridges, H.R., J.N. Blaza, Z. Yin, I. Chung, M.N. Pollak, and J. Hirst, Structural basis of mammalian respiratory complex I inhibition by medicinal biguanides. Science, 2023. 379(6630): p. 351-357, 10.1126/science.ade3332.

[34] Grba, D.N., I. Chung, H.R. Bridges, A.-N.A. Agip, and J. Hirst, Investigation of hydrated channels and proton pathways in a high-resolution cryo-EM structure of mammalian complex I. Science Advances, 2023. 9(31): p. eadi1359, 10.1126/sciadv.adi1359.

[35] Kampjut, D. and L.A. Sazanov, The coupling mechanism of mammalian respiratory complex I. Science, 2020. 370(6516): p. eabc4209, 10.1126/science.abc4209.

[36] Narayanan, M., J.A. Sakyiama, M.M. Elguindy, and E. Nakamaru-Ogiso, Roles of subunit NuoL in the proton pumping coupling mechanism of NADH: ubiquinone oxidoreductase (complex I) from Escherichia coli. The journal of biochemistry, 2016. 160(4): p. 205–215, 10.1093/jb/mvw027.

[37] Michel, J., J. DeLeon-Rangel, S. Zhu, K. Van Ree, and S.B. Vik, Mutagenesis of the L, M, and N subunits of complex I from Escherichia coli indicates a common role in function. PloS one, 2011. 6(2): p. e17420, 10.1371/journal.pone.0017420.

[38] Torres-Bacete, J., E. Nakamaru-Ogiso, A. Matsuno-Yagi, and T. Yagi, Characterization of the NuoM (ND4) subunit in Escherichia coli NDH-1: conserved charged residues essential for energy-coupled activities. Journal of Biological Chemistry, 2007. 282(51): p. 36914–36922, 10.1074/jbc.M707855200.

[39] Sato, M., P.K. Sinha, J. Torres-Bacete, A. Matsuno-Yagi, and T. Yagi, Energy transducing roles of antiporter-like subunits in Escherichia coli NDH-1 with main focus on subunit NuoN (ND2). Journal of Biological Chemistry, 2013. 288(34): p. 24705–24716, 10.1074/jbc.M113.482968.

[40] Röpke, M., P. Saura, D. Riepl, M.C. Pöverlein, and V.R. Kaila, Functional water wires catalyze long-range proton pumping in the mammalian respiratory complex I. Journal of the American Chemical Society, 2020. 142(52): p. 21758–21766, 10.1021/jacs.0c09209.

[41] Di Luca, A., A.P. Gamiz-Hernandez, and V.R. Kaila, Symmetry-related proton transfer pathways in respiratory complex I. Proceedings of the National Academy of Sciences, 2017. 114(31): p. E6314–E6321, 10.1073/pnas.1706278114.

[42] Haapanen, O. and V. Sharma, Role of water and protein dynamics in proton pumping by respiratory complex I. Scientific reports, 2017. 7(1): p. 7747, 10.1038/s41598-017-07930-1.

[43] Tan, P., Z. Feng, L. Zhang, T. Hou, and Y. Li, The mechanism of proton translocation in respiratory complex I from molecular dynamics. Journal of Receptors and Signal Transduction, 2015. 35(2): p. 170–179, 10.3109/10799893.2014.942464.

[44] Mathiesen, C. and C. Hägerhäll, Transmembrane topology of the NuoL, M and N subunits of NADH: quinone oxidoreductase and their homologues among membrane-bound hydrogenases and bona fide antiporters. Biochimica et Biophysica Acta (BBA)- Bioenergetics, 2002. 1556(2-3): p. 121–132, 10.1016/S0005-2728(02)00343-2.

[45] Zhuang, J., J.H. Amoroso, R. Kinloch, J.H. Dawson, M.J. Baldwin, and B.R. Gibney, Evaluation of electron-withdrawing group effects on heme binding in designed proteins: implications for heme a in cytochrome c oxidase. Inorganic Chemistry, 2006. 45: p. 4685–4694, 10.1021/ic060072c.

[46] Fontecilla-Camps, J.C., A. Volbeda, C. Cavazza, and Y. Nicolet, Structure/function relationships of [NiFe]-and [FeFe]-hydrogenases. Chemical reviews, 2007. 107(10): p. 4273–4303, 10.1021/cr050195z.

[47] Efremov, R.G. and L.A. Sazanov, The coupling mechanism of respiratory complex I—a structural and evolutionary perspective. Biochimica et Biophysica Acta (BBA)- Bioenergetics, 2012. 1817(10): p. 1785–1795, 10.1016/j.bbabio.2012.02.015.

[48] Kaila, V.R., Resolving chemical dynamics in biological energy conversion: Long-range proton-coupled electron transfer in respiratory complex I. Accounts of Chemical Research, 2021. 54(24): p. 4462–4473, 10.1021/acs.accounts.1c00524.

[49] Sazanov, L.A., The mechanism of coupling between electron transfer and proton translocation in respiratory complex I. Journal of bioenergetics and biomembranes, 2014. 46: p. 247–253, 10.1007/s10863-014-9554-z.

[50] Verkhovskaya, M. and D.A. Bloch, Energy-converting respiratory Complex I: on the way to the molecular mechanism of the proton pump. The international journal of biochemistry & cell biology, 2013. 45(2): p. 491–511, 10.1016/j.biocel.2012.08.024.

[51] Gupta, C., U. Khaniya, C.K. Chan, F. Dehez, M. Shekhar, M. Gunner, L. Sazanov, C. Chipot, and A. Singharoy, Charge transfer and chemo-mechanical coupling in respiratory complex I. Journal of the American Chemical Society, 2020. 142(20): p. 9220–9230, 10.1021/jacs.9b13450.

[52] Gutiérrez-Fernández, J., K. Kaszuba, G.S. Minhas, R. Baradaran, M. Tambalo, D.T. Gallagher, and L.A. Sazanov, Key role of quinone in the mechanism of respiratory complex I. Nature communications, 2020. 11(1): p. 4135, 10.1038/s41467-020-17957-0.

[53] Gupta, C., U. Khaniya, C.K. Chan, M. Gunner, C. Chipot, F. Dehez, and A. Singharoy, Chemomechanical Coupling of Mitochondrial Complex I. Biophysical Journal, 2019. 116(3): p. 155a, 10.1016/j.bpj.2018.11.858.

[54] Sharma, V., G. Belevich, A.P. Gamiz-Hernandez, T. Róg, I. Vattulainen, M.L. Verkhovskaya, M. Wikström, G. Hummer, and V.R. Kaila, Redox-induced activation of the proton pump in the respiratory complex I. Proceedings of the National Academy of Sciences, 2015. 112(37): p. 11571–11576, 10.1073/pnas.150376111.

[55] Wei, R.J., Y. Zhang, J. Mao, D. Kaur, U. Khaniya, and M. Gunner, Comparison of proton transfer paths to the QA and QB sites of the Rb. sphaeroides photosynthetic reaction centers. Photosynthesis Research, 2022. 152(2): p. 153–165, 10.1007/s11120-022-00906-x.

[56] Khaniya, U., J. Mao, R.J. Wei, and M. Gunner, Characterizing protein protonation microstates using Monte Carlo sampling. The Journal of Physical Chemistry B, 2022. 126(13): p. 2476–2485, 10.1021/acs.jpcb.2c00139.

[57] Song, Y., J. Mao, and M.R. Gunner, MCCE2: improving protein pKa calculations with extensive side chain rotamer sampling. Journal of computational chemistry, 2009. 30(14): p. 2231–2247, 10.1002/jcc.21222.

[58] Song, Y., J. Mao, and M.R. Gunner, Calculation of proton transfers in bacteriorhodopsin bR and M intermediates. Biochemistry, 2003. 42(33): p. 9875–9888, 10.1021/bi034482d.

[59] Gowers, R.J., M. Linke, J. Barnoud, T.J.E. Reddy, M.N. Melo, S.L. Seyler, J. Domanski, D.L. Dotson, S. Buchoux, and I.M. Kenney. MDAnalysis: a Python package for the rapid analysis of molecular dynamics simulations. 2019. Los Alamos National Laboratory (LANL), Los Alamos, NM (United States), 10.25080/Majora-629e541a-00e.

[60] Michaud-Agrawal, N., E.J. Denning, T.B. Woolf, and O. Beckstein, MDAnalysis: a toolkit for the analysis of molecular dynamics simulations. Journal of computational chemistry, 2011. 32(10): p. 2319–2327, 10.1002/jcc.21787.

[61] Tubiana, T., J.-C. Carvaillo, Y. Boulard, and S. Bressanelli, TTClust: a versatile molecular simulation trajectory clustering program with graphical summaries. Journal of chemical information and modeling, 2018. 58(11): p. 2178–2182, 10.1021/acs.jcim.8b00512.

[62] Gunner, M., X. Zhu, and M.C. Klein, MCCE analysis of the pKas of introduced buried acids and bases in staphylococcal nuclease. Proteins: Structure, Function, and Bioinformatics, 2011. 79(12): p. 3306–3319, 10.1002/prot.23124.

[63] Cai, X., K. Haider, J. Lu, S. Radic, C.Y. Son, Q. Cui, and M. Gunner, Network analysis of a proposed exit pathway for protons to the P-side of cytochrome c oxidase. Biochimica et Biophysica Acta (BBA)-Bioenergetics, 2018. 1859(10): p. 997–1005, 10.1016/j.bbabio.2018.05.010.

[64] Mühlbauer, M.E., P. Saura, F. Nuber, A. Di Luca, T. Friedrich, and V.R. Kaila, Water-gated proton transfer dynamics in respiratory complex I. Journal of the American Chemical Society, 2020. 142(32): p. 13718–13728, 10.1021/jacs.0c02789.

[65] Tocilescu, M.A., U. Fendel, K. Zwicker, S. Dröse, S. Kerscher, and U. Brandt, The role of a conserved tyrosine in the 49-kDa subunit of complex I for ubiquinone binding and reduction. Biochimica et Biophysica Acta (BBA)-Bioenergetics, 2010. 1797(6-7): p. 625–632, 10.1016/j.bbabio.2010.01.029.

[66] Tocilescu, M.A., U. Fendel, K. Zwicker, S. Kerscher, and U. Brandt, Exploring the ubiquinone binding cavity of respiratory complex I. Journal of Biological Chemistry, 2007. 282(40): p. 29514–29520, 10.1074/jbc.M704519200.

[67] Gamiz-Hernandez, A.P., A. Jussupow, M.P. Johansson, and V.R. Kaila, Terminal electron–proton transfer dynamics in the quinone reduction of respiratory complex I. Journal of the American Chemical Society, 2017. 139(45): p. 16282–16288, 10.1021/jacs.7b08486.

[68] Sazanov, L.A., Respiratory complex I: mechanistic and structural insights provided by the crystal structure of the hydrophilic domain. Biochemistry, 2007. 46(9): p. 2275–2288, 10.1021/bi602508x.

[69] Efremov, R.G. and L.A. Sazanov, Structure of the membrane domain of respiratory complex I. Nature, 2011. 476(7361): p. 414-420, 10.1038/nature10330.

[70] Sinha, P.K., J. Torres-Bacete, E. Nakamaru-Ogiso, N. Castro-Guerrero, A. Matsuno-Yagi, and T. Yagi, Critical roles of subunit NuoH (ND1) in the assembly of peripheral subunits with the membrane domain of Escherichia coli NDH-1. Journal of biological chemistry, 2009. 284(15): p. 9814–9823, 10.1074/jbc.M809468200.

[71] Lu, J. and M. Gunner, Characterizing the proton loading site in cytochrome c oxidase. Proceedings of the National Academy of Sciences, 2014. 111(34): p. 12414–12419, 10.1073/pnas.1407187111.

[72] Letts, J.A., K. Fiedorczuk, G. Degliesposti, M. Skehel, and L.A. Sazanov, Structures of respiratory supercomplex I+ III2 reveal functional and conformational crosstalk. Molecular cell, 2019. 75(6): p. 1131–1146. e6, 10.1016/j.molcel.2019.07.022.

[73] Steimle, S., M. Willistein, P. Hegger, M. Janoschke, H. Erhardt, and T. Friedrich, Asp563 of the horizontal helix of subunit NuoL is involved in proton translocation by the respiratory complex I. FEBS letters, 2012. 586(6): p. 699–704, 10.1016/j.febslet.2012.01.056.

[74] Euro, L., G. Belevich, M.I. Verkhovsky, M. Wikström, and M. Verkhovskaya, Conserved lysine residues of the membrane subunit NuoM are involved in energy conversion by the proton-pumping NADH: ubiquinone oxidoreductase (Complex I). Biochimica et Biophysica Acta (BBA)-Bioenergetics, 2008. 1777(9): p. 1166–1172, 10.1016/j.bbabio.2008.06.001.

[75] Torres-Bacete, J., P.K. Sinha, A. Matsuno-Yagi, and T. Yagi, Structural contribution of C-terminal segments of NuoL (ND5) and NuoM (ND4) subunits of complex I from Escherichia coli. Journal of Biological Chemistry, 2011. 286(39): p. 34007–34014, 10.1074/jbc.M111.260968.

[76] Kao, M.-C., S. Di Bernardo, M. Perego, E. Nakamaru-Ogiso, A. Matsuno-Yagi, and T. Yagi, Functional roles of four conserved charged residues in the membrane domain subunit NuoA of the proton-translocating NADH-quinone oxidoreductase from Escherichia coli. Journal of Biological Chemistry, 2004. 279(31): p. 32360–32366, 10.1074/jbc.M403885200.

[77] Kurki, S., V. Zickermann, M. Kervinen, I. Hassinen, and M. Finel, Mutagenesis of three conserved Glu residues in a bacterial homologue of the ND1 subunit of complex I affects ubiquinone reduction kinetics but not inhibition by dicyclohexylcarbodiimide. Biochemistry, 2000. 39(44): p. 13496–13502, 10.1021/bi001134s.

[78] Belevich, G., L. Euro, M. Wikström, and M. Verkhovskaya, Role of the conserved arginine 274 and histidine 224 and 228 residues in the NuoCD subunit of complex I from Escherichia coli. Biochemistry, 2007. 46(2): p. 526–533, 10.1021/bi062062t.

[79] Crooks, G.E., G. Hon, J.-M. Chandonia, and S.E. Brenner, WebLogo: a sequence logo generator. Genome research, 2004. 14(6): p. 1188–1190, http://www.genome.org/cgi/doi/10.1101/gr.849004.

[80] Gu, J., T. Liu, R. Guo, L. Zhang, and M. Yang, The coupling mechanism of mammalian mitochondrial complex I. Nature Structural & Molecular Biology, 2022. 29(2): p. 172–182, 10.1038/s41594-022-00722-w.

[81] Stuchebrukhov, A.A. and T. Hayashi, Single protonation of the reduced quinone in respiratory complex I drives four-proton pumping. FEBS letters, 2023. 597(2): p. 237–245, 10.1002/1873-3468.14518.

[82] Sazanov, L.A., From the ‘black box’to ‘domino effect’mechanism: what have we learned from the structures of respiratory complex I. Biochemical Journal, 2023. 480(5): p. 319–333, 10.1042/BCJ20210285.

[83] Zdorevskyi, O., A. Djurabekova, J. Lasham, and V. Sharma, Horizontal proton transfer across the antiporter-like subunits in mitochondrial respiratory complex I. Chemical Science, 2023. 14(23): p. 6309–6318, 10.1039/D3SC01427D.

[84] Kravchuk, V., O. Petrova, D. Kampjut, A. Wojciechowska-Bason, Z. Breese, and L. Sazanov, A universal coupling mechanism of respiratory complex I. Nature, 2022. 609(7928): p. 808-814, 10.1038/s41586-022-05199-7.

[85] Sazanov, L.A. and P. Hinchliffe, Structure of the hydrophilic domain of respiratory complex I from Thermus thermophilus. science, 2006. 311(5766): p. 1430-1436, 10.1126/science.1123809.

