## Supplemental Information for "Finding the E-Channel Proton Loading Sites by Calculating the Ensemble of Protonation Microstates"

**SI.1.** Methods
**SI.1.1.** Molecular Dynamics simulation

**SI.1.2.** IPECE membrane creation

**SI.1.3.** MCCE force field

**SI.1.4.** Monte Carlos sampling

**SI.1.5.** Correlation of residue protonation in MC microstates **SI.1.6.** Correlation study

**SI.1.6.** Multiple sequence analysis

**Figure SI.1.** Percentages of the frames isolated as loaded and unloaded and mixed loaded-unloaded frames

**Figure SI.2. Distribution of unique protonation microstates for the (a) Apo and (b) MQH2-docked snapshots.**

**Figure SI.3.** Correlation of Central Cluster residues protonation in loaded, and unloaded, frames.

**Figure SI.4.** Correlation studies of the N-side cluster residues

**Figure SI.5.** Correlation between the Central Cluster and Cluster-5 total charge

**Figure SI.6.** Movement of the residues 4-His38 and 4-Asp139 in the Q region

**Figure SI.7.** Proton pumping cycle

**Table SI.1.** Nomenclature of the core subunits of complex I in different organism

**Table SI.2.** All the residues of the Central Cluster and the N-side cluster

**Table SI.3.** Average protonation microstates in ten loaded and ten unloaded snapshots.

**Table SI.4.** Highest probable protonation microstates found in MCCE analysis of snapshots where the Central Cluster is loaded (top), mix loaded-unloaded (middle) and unloaded (bottom)

**Table SI.5.** A summary table of experimental side-directed mutagenesis studies that aligned with most of our computationally predicted active residues in Central Cluster and N-side cluster PLS residues and the covariance data.

**Table SI.6.** List of the residues with the residues of within 5.0 Å of the active residues in the Central Cluster and the N-side cluster.

**SI.1. Methos
SI.1.1. Molecular Dynamics simulation**

*Thermus thermophilus* Complex I, is immersed in a hydrated patch of 1000 1-palmitoyl-2-oleoyl-sn-glycero-3-phosphocholines (POPC) in the presence of 150 mM NaCl. The system has ~1,000,000 atoms in a 28 nm x 14 nm x 24 nm box [1, 2]. Complex I subunits Nqo4, 7, 8, 12-14, Asp, Glu, Arg, and Lys are ionized, while His are neutral with the proton on ND1. The protonation state of all other titratable residues in other different subunits was determined using pK_a_ values estimated by PropK_a_ [1]. The CHARMM36 force field, incorporating CMAP corrections for proteins [3, 4] and the TIP3P model for water is used. The parameters for Iron-sulfur clusters are taken from Density Functional Theory calculations by Chang and Kim [5] Iron-sulfur clusters N1a, N1b, N5, N7, and N6b ware oxidized while N3, N4, N6a, and N2, reduced. This is based on the pattern supported on bovine Complex I [6-8].The parameter set for Flavin mononucleotide are from Freddolino et. al. [9].

The system undergoes equilibration in the NPT ensemble (temperature T = 313 K, pressure P = 1 atm) for100 nanoseconds in NAMD. The production run uses 1.0 fs timesteps, with a force-based switching function for long-range interactions within the range of 10 to 12 Å. A Langevin thermostat, Nosé-Hoover Langevin barostat is used, with a flexible cell. Trajectories are generated for a total duration of 0.5 microsecond [2].

**SI.1.2. IPECE membrane creation for MCCE calculations**

To create a low dielectric region to surround the five E-channel peptides, IPECE [10] is run with only the membrane embedded peptides, Nqo7, 8, 10, and 11. It orients the protein and inserts it into a 30 Å wide, low dielectric region. Nqo4, in the puerperal arm, is then returned to complete the structure.

**SI.1.3. MCCE energy**

A microstate in the protein is one charge/position conformer for each residue and ligand [10]. The microstate energy is composed of reference energy, self and pairwise terms [10, 11]. The energy (∆H^x^) of microstate x is expressed as:

$$\Delta H^{x}=\sum_{i=1}^{M} \delta_{x,i}\{\left[ 2.3m_{i}k_{b}T\left( pH -\mathrm{pK}_{sol,i} \right)+n_{i}F\left( E_{h} -E_{m sol,i} \right) \right]$$

${+ \Delta\Delta G}_{rxn,i}+{\Delta\Delta G}_{bkbn,i}^{CE}+{\Delta\Delta G}_{bkbn,i}^{LJ}{+ \Delta\Delta G}_{torsion,i}+$ ${\Delta\Delta G}_{SAS,i}$

$$+\sum_{j=i+1}^{M} \delta_{x,j}[{\Delta G}_{ij}^{CE}+{\Delta G}_{ij}^{LJ}]\} (S1)$$

The first line in equation S1 details the energy each residue would have in solution where pK_a_=pK_a,sol,_, given its microstate assigned protonation state. M represents the total number of conformers, δ_x,i_ is 1 if conformer i is present in microstate x, or 0 otherwise. m_i_ is 1 for basic protonated, -1 for acidic deprotonated, and 0 for neutral conformers. pK_sol,i_ is the reference pK_a_ of the residue type in solution, E_m_ is the reference electrochemical midpoint potential of the redox-active group. F denotes the Faraday constant, n_i_ is the change in the number of electrons on a conformer participating in a redox titration, while pH and E_h_ are relevant solution parameters describing the chemical potential of protons or electrons in equilibrium with the protein.

The second line finds the self-energies of the conformer forming the microstates. These energies, independent of conformer choices for other residues, include the loss of conformer solvation energy during its transition from solution to its position in the protein, torsion energy, continuum electrostatic (CE), and Lennard Jones (LJ) van der Waals interactions with fixed backbone amides. Additionally, it adds the favorable van der Waals interactions of exposed side chain surface with the implicit solvent [10, 12].

The third line adds the continuum electrostatic and Lennard Jones pairwise interactions between each pair of conformers in different residues. MCCE has been tested against available experimental pK_a_s [13, 14] and E_m_s [15].

All energies are computed before Monte Carlo (MC) sampling. Electrostatic energies are determined using the Poisson-Boltzmann solver in Delphi [15-17], employing Parse charges [18], a dielectric constant of 4 for the protein, 80 for the solvent, and an implicit salt concentration of 150 mM. MCCE adjusts continuum electrostatic energies for changes in the dielectric boundary due to alterations in surface rotamers [10]. Non-electrostatic energies utilize the Amber force field [19].

**SI.1.4. Monte Carlos sampling**

In MCCE (MultiConformation Continuum Electrostatics), transitions between microstates occur through stochastic selections. Initially, a residue is chosen, followed by the selection of an available conformer for that residue. If the selected residue contains conformers interacting with an absolute energye greater than 0.5 kcal/mol with a conformer of another residue, there is a 50% chance of additional conformer changes for that other residue. This process involves swapping conformers in multiple residues, and a decision is made using the Metropolis–Hasting’s algorithm to either accept or reject the change(s). Allowing one to three residues to change helps prevent clashes between strongly coupled groups and aids convergence [20-22].

The Monte Carlo sampling procedure starts with a random microstate, assigning one conformer to each residue. Then the temperature is gradually reduced in the Metropolis-Hastings acceptance criteria. Following annealing, conformer reduction is applied, and then MC sampling is performed at room temperature. Rarely chosen conformers (probability < 0.001) are removed from the list before the production phase to enhance convergence, targeting an acceptance rate of approximately 30%. Early removal of conformers may be present in higher energy states, and adjustments can be modified to enhance speed or to not miss rare states [23]. The discarded conformers are excluded from all subsequent sampling. Residues with a probability of a single conformer being unity are fixed residues and are removed from the sampling list. All remaining free residues retain the choice of conformers. Residues with a fixed protonation state can still sample conformers with different positions and remain on the free residue list.

**SI.1.5. Correlation of residue protonation in MC microstates**

A correlation coefficient is used to assess the linear association between a pair of variables. The value lies between -1 to +1. A negative correlation coefficient reveals an inverse relationship, so as one variable increases, the other tends to decrease. Here that will indicate when one group is ionized its partner tends to be neutral. Correlation values near ±1 indicate a robust and consistent relationship, while coefficients approaching 0 signify a lack of connection.

Knowing individual microstates allows us to identify when residue protonation states are coupled together. Correlation is evaluated using the weighted Pearson Correlation coefficient (𝑟) as described previously [24].

$$\mathbf{r}_{\mathbf{pq}}\mathbf{=}\frac{\sum_{\mathbf{i=1}}^{\mathbf{n}} \mathbf{w}_{\mathbf{i}}\mathbf{(}\mathbf{p}_{\mathbf{i}}\mathbf{-}\bar{\mathbf{p}}\mathbf{)(}\mathbf{q}_{\mathbf{i}}\mathbf{-}\bar{\mathbf{q}}\mathbf{)}}{\sqrt{\sum_{\mathbf{i=1}}^{\mathbf{n}} \mathbf{(}\mathbf{w}_{\mathbf{i}}\left( \mathbf{p}_{\mathbf{i}}\mathbf{-}\bar{\mathbf{p}} \right)^{\mathbf{2}}\mathbf{)}\sum_{\mathbf{i=1}}^{\mathbf{n}} \mathbf{(}\mathbf{w}_{\mathbf{i}}{\mathbf{(}\mathbf{q}_{\mathbf{i}}\mathbf{-}\bar{\mathbf{q}}\mathbf{)}}^{\mathbf{2}}}}\mathbf{(}S2\mathbf{)}$$

r_pq_ is the correlation between residue p and q, p_i_ and q_i_ are the protonation states for two residues in a microstate, i, 𝑝̅ and $\bar{q}$ are the mean value of p_i_ and q_i_ respectively and n is the number of unique accepted protonation microstates. The sum runs over unique protonation microstates, so the weight w_i_, is equal to the number of times this state is found in the ensemble.

**SI.1.6. Multiple sequence analysis**

Multiple sequence alignment was carried out on 1000 Complex I sequences using NCBI Blast [25], and Clustal Omega [25]. Aligned residues of interest were extracted as input for Weblogo [26].

FIGURES

**Figure SI.1. Distribution of unique protonation microstates representative of the (a) Apo and (b) MQH2-docked snapshots.**

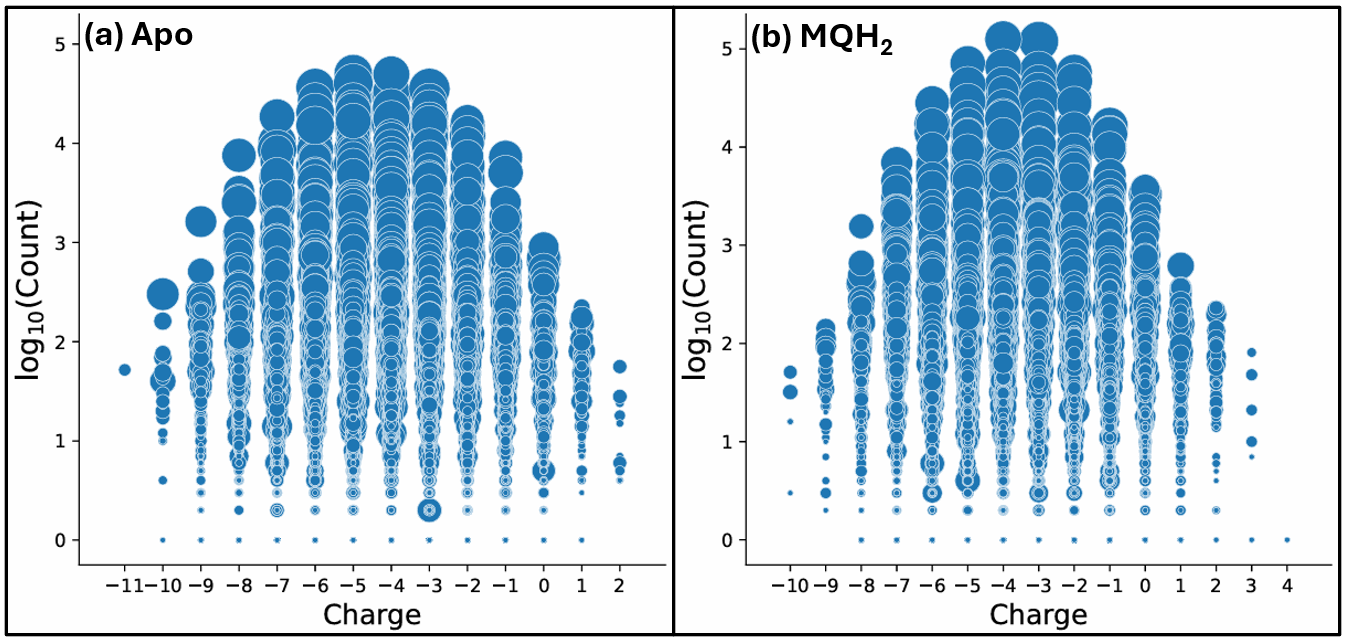

**(a)** Distribution of unique protonation microstates **of** five E-channel subunits of apo snapshoot, average ensemble charge of -4.3, charges ranging from -11 to +2 with the most probable charges between -6 and -3, centered between -5 to -4 and >26,000 accepted protonation microstates, and **(b)** MQH_2_-docked snapshoot, average ensemble charge of -3.4 charges ranging from -10 to +4 with the most probable charges between -5 and -2, centered between -4 to -3 and >32,000 protonation microstates. Each shows the probability in log_10_(count) and summed charge of one protonation microstate. Dots in a column are protein tautomers with different locations of the protons, but the same total charge. Dot size indicates the range of energies of the conformational microstates that are found in this protonation state.

**Figure SI.2. Percentage of frames identified with the Central and N-side clusters classified as loaded, unloaded or mixed loaded-unloaded from the apo, MQ, and MQH_2_ Complex I MD trajectories.**

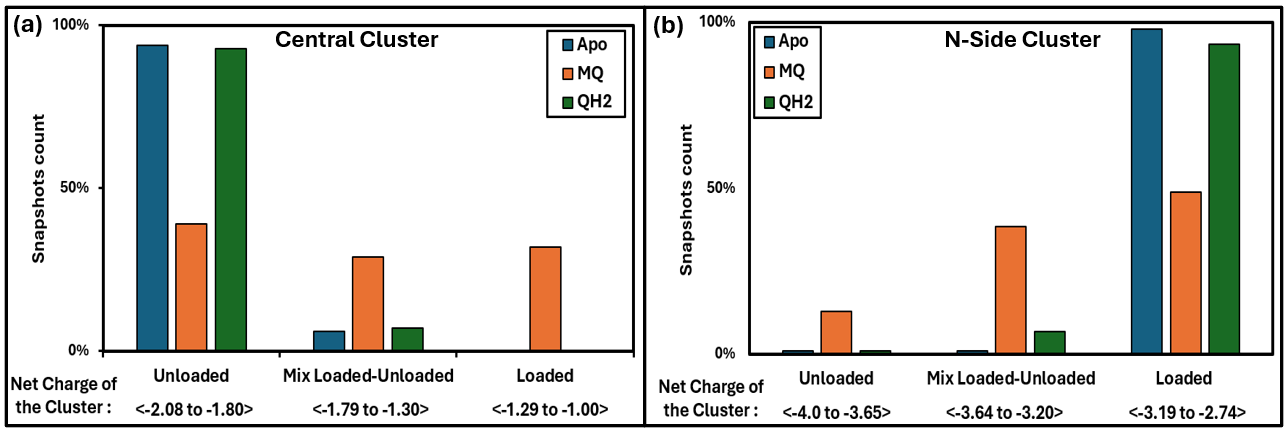

Blue: 20 apo snapshots; red: 40 MQ snapshots; green: 20 MQH_2_ snapshot**s (a) Central Cluster:** Average summed charge of residues 8-E213, 8-H251, 8-E130, 8-E163, 7-D72, and 7-E74 in MC sampling of a snapshot. Snapshots with Central Cluster net charge from -2.08 to -1.80 are unloaded; -1.29 to -1.00 are loaded and those with a net charge from -1.79 to -1.30 are partially loaded. **(b) N-side cluster:** Average summed charge of residues 8-E225, 4-E51, 4-E50, 7-H60, and 4-E216. Snapshots with N-side cluster net charge from -4.08 to -3.65 are unloaded; -3.64 to -3.20 are loaded and those with a net charge from -3.19 to -2.74 are partially loaded.

**Figure SI.3. Correlation of Central Cluster residue protonation in loaded, and unloaded, frames.**

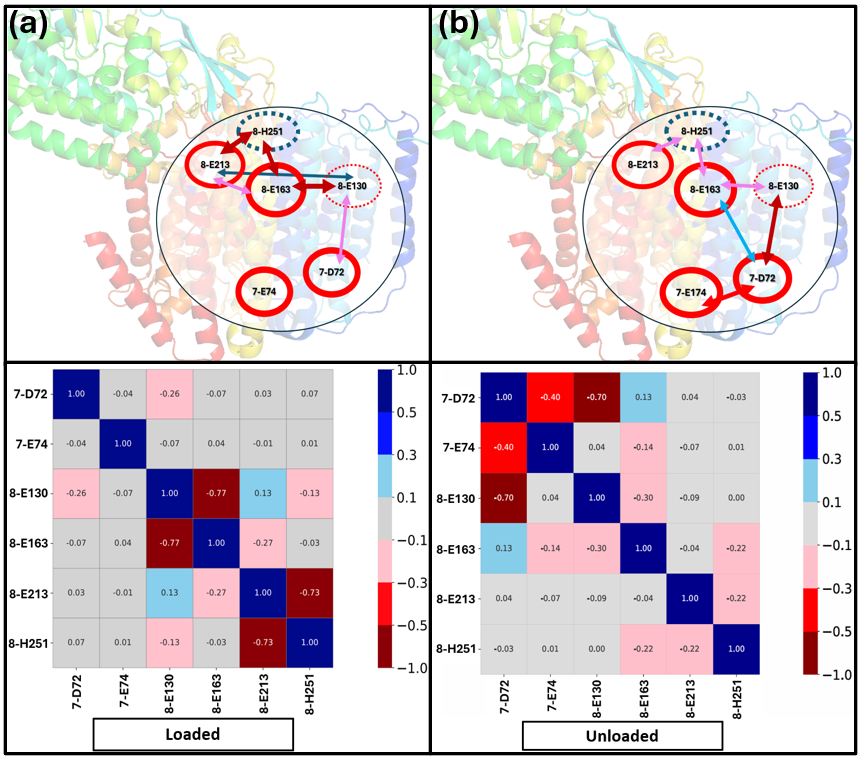

Snapshot with loaded Central Cluster; average charge (a) -1.10; (b) -1.99. (Top) The Central Cluster residues whose protonation states change between the high probability loaded and unloaded microstates. The arrows indicate the coupling between the residues (bottom). Red arrow: Negatively correlated protonation changes (one residue is more ionized and the other less); Blue arrows: positive correlation. Arrow thickness indicates correlation strength. Circles around residue number reflect residue conservation in a multiple sequence alignment (Fig. 8). Solid circles: highly conserved; dashed circles: lower conservation. Red circles: acids; Blue: bases. (Bottom) The Pearson weighted correlation coefficient (Equation 1). Only residues exhibiting an absolute correlation value of ≥0.1 with at least one other residue are included. Each square reports the correlation strength between two residues. The color spectrum denotes correlation values: dark blue (0.5 to 1.0), blue (0.3 to 0.5), sky blue (0.1 to 0.3), light gray (−0.1 to 0.1), orange (−0.3 to −0.1), red (−0.5 to −0.3), and dark red (−1.0 to −0.5). The correlations shown here are the source of the weight and color of the arrows in the top panel.

**Figure SI.4. Correlation studies of the N-side cluster residues.**

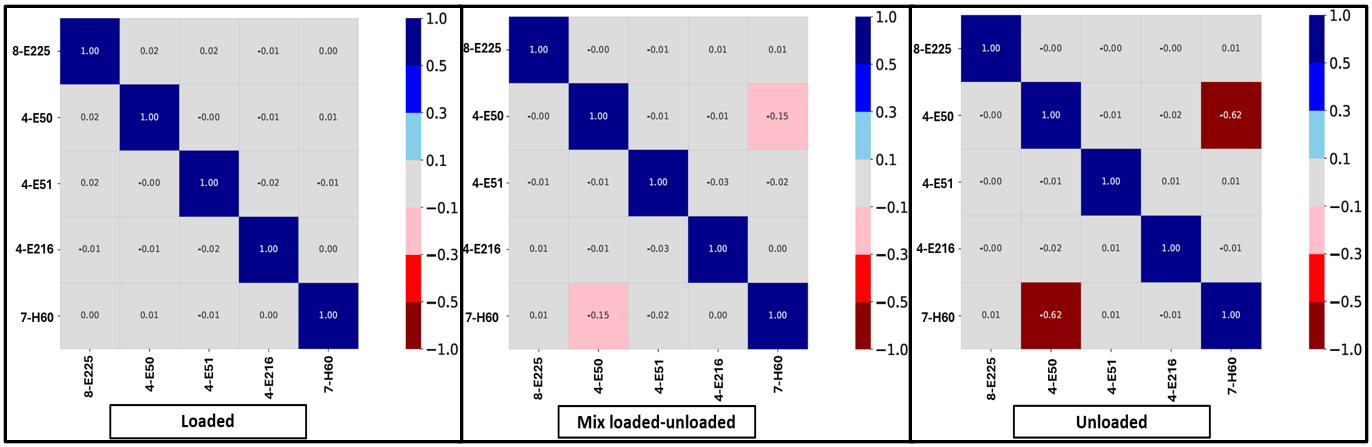

The Pearson's weighted correlation coefficient is shown and only residues exhibiting an absolute correlation value of ≥0.1 with at least one other residue are included. See Fig. SI.2 for more complete description. In unloaded snapshots there is a strong negative correlation between 7-H60 and 4-E50, which are 6.3 Å apart, indicating when one His is protonated the other is less likely to be. In the loaded state no correlation is seen because both residues are always neutral.

**Figure SI.5. Lack of correlation between the Central Cluster and Cluster 5 [2] total charge in Apo-, MQ and MQH_2_ docked trajectories.**

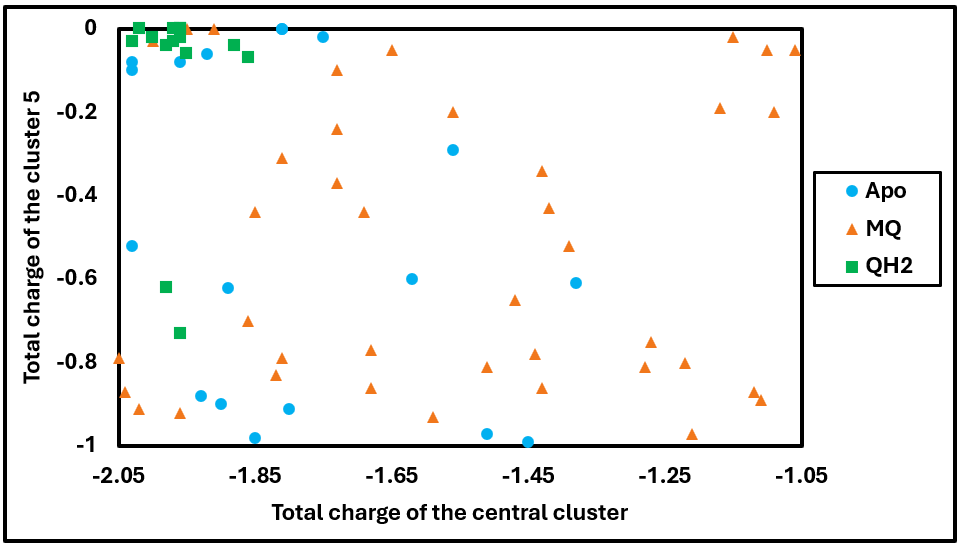

Each data point represents the MCCE average charge in a snapshot. The color and shapes indicate snapshots from MQ (orange triangle), MQH_2_ (green square), and apo (blue circle) trajectories. The charges of the two clusters are not correlated. In the MQH_2_ docked trajectory the Central Cluster is mostly unloaded while Cluster 5 is mostly protonated. The residues in Cluster 5 are 10-Y43, 10-Q55, 10-Y59, and 11-E32.

**Figure SI.6. Movement of 4-His38 ligand and nearby 4-Asp139.**

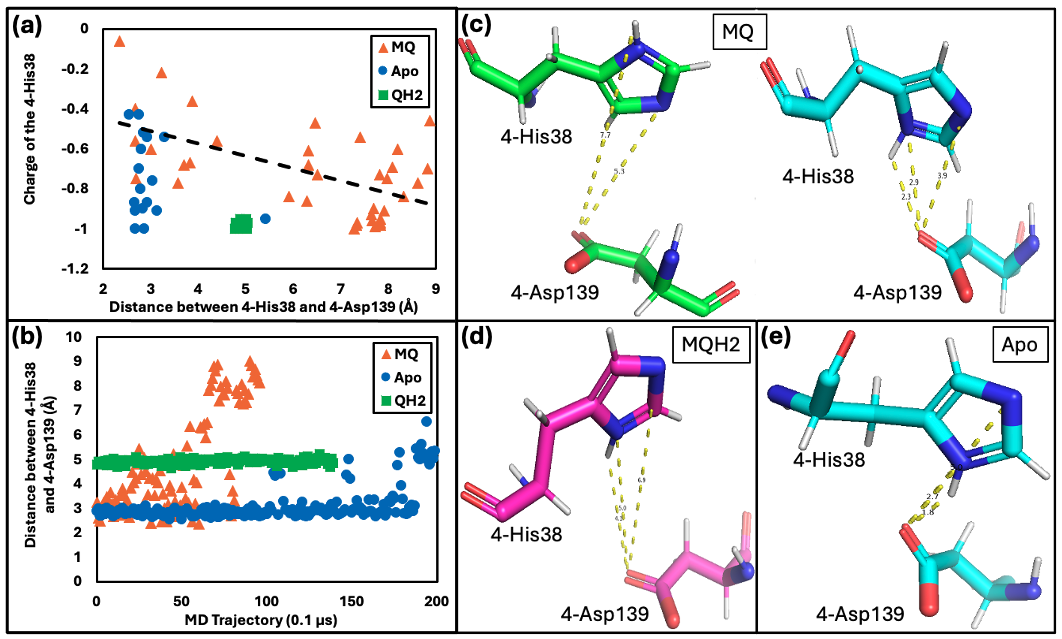
The residue 4-H38 is a ligand to MQ and can make a hydrogen bond to 4-D139. **(a)** Correlation of sum charge of His + Asp with the distance between the two residues. Asp is always ionized. Each data point is the average charge in MC sampling for one snapshot. All MQH_2_ data overlaps. **(b)** Distance between 4-H38 ND1 and 4-E139 OE1 in the MQ (orange triangle), MQH_2_ (green square), and apo (blue circle) trajectories. **(c)** Position of 4-H38 and 4-D139 in representative MQ-docked snapshots; **left**: neutral 4-His38 (average charge: 0.03) and **right**: 4-His38 is ionized (+1).4-D139 is always ionized. **(d, e)** Position of 4-H38 and 4-D139 in MQH_2_, and apo-snapshots.

**Figure SI.7. Proton pumping cycle.**

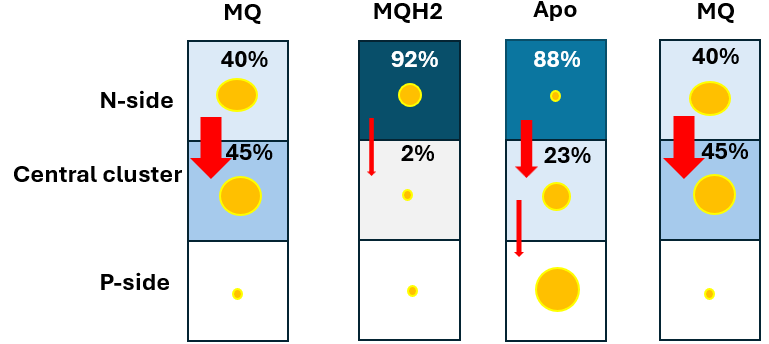

Schematic of proton pumping cycle illustrating hydrogen bond connectivity and PLS loading in snapshots from MQ, MQH_2_, and apo trajectories. Darker squares are more likely to be loaded. The percentage loaded are given in the boxes. The size of the yellow circle is proportional to the degree of H-bond connectivity of the cluster residue found previously by Khaniya et. al. [2]; and the size of the red arrows is proportional to the inter cluster connectivity. Larger circles show more intracluster connections; larger arrows indicate more connections between clusters.

It is not known how pumping is coupled to quinone reduction. Between MQ and MQH_2_ the quinone is reduced and protonated twice. The system could feel the charge of an anionic semiquinone (MQ^-^) or quinol (MQH^-^) intermediates. Alternatively, a quinone could bind a proton each time it is reduced, remaining neutral, but changing the hydrogen bond pattern around the quinone binding site. Alternative binding sites for the exiting quinone may also play a role [27, 28]. An unprotonated, doubly reduced species would be expected to be highly unfavorable within the protein environment [29].

**TABLES**

**Table SI.1. Nomenclature of the core subunits of complex I assigned in different organism**

| Region | *Thermus thermophilus* [30] *Paracoccus denitirificans* [31] | *Escherichia coli* [30] | *Bos taurus* [30] | *Yarrowia lipolytica* [31] | *Homo sapiens* [30] [*Mus musculus*](https://www.rcsb.org/search?q=rcsb_entity_source_organism.taxonomy_lineage.name:Mus%20musculus) [32] |
| --- | --- | --- | --- | --- | --- |
| Peripheral arm | Nqo1 | NuoF | 51kDa | NUBM | NDUFV1 |
|  | Nqo2 | NuoE | 24kDa | NUHM | NDUFV2 |
|  | Nqo3 | NuoG | 75kDa | NUAM | NDUFS1 |
|  | Nqo4 | NuoD | 49kDa | NUCM | NDUFS2 |
|  | Nqo5 | NuoC | 30kDa | NUGM | NDUFS3 |
|  | Nqo6 | NuoB | PSST | NUKM | NDUFS7 |
|  | Nqo9 | NuoI | TYKY | NUIM | NDUFS8 |
| Membrane arm | Nqo7 | NuoA | ND3 | NU3M | ND3 |
|  | Nqo8 | NuoH | ND1 | NU1M | ND1 |
|  | Nqo10 | NuoJ | ND6 | NU6M | ND6 |
|  | Nqo11 | NuoK | ND4L | NULM | ND4L |
|  | Nqo12 | NuoL | ND5 | NU5M | ND5 |
|  | Nqo13 | NuoM | ND4 | NU4M | ND4 |
|  | Nqo14 | NuoN | ND2 | NU2M | ND2 |

Unfortunately, the same subunits are assigned different names in Complex I from different species. The chain designations for *Thermus thermophilus* are used here.

**Table SI.2. All residues previously identified in the Central and the N-side clusters [2].** The residues from Nqo4, 7, and 8 (PDB ID: 4HEA chains E, P, and Q) of *T. thermophilus* complex I involved in Central Cluster (cluster 4 in ref [2]) and N-side Cluster (cluster 1, 2, and 3 in [2]). The red highlighted residues change protonation state in different microstates reported here. The underlined residues are on the surface exposed, and the bold residues are in the peripheral arm.

| Cluster name | Residues name (protonatable resides**;** others) | *T. thermophilus* Subunits  (4HEA chain ID) |
| --- | --- | --- |
| Central Cluster | D72, E74, **E110**, **K113;** T10, W79, T100, W111, **Y109**, **R117** | Nqo7 (P) |
|  | E163, E130, E213, H251**;** S89, Y124, S129, Y134, S161, Y162, Y206, S210, T254, W310 | Nqo8 (Q) |
|  | **E158; T150** | Nqo10 (R) |
| N-side cluster | **E51, H58**, **E216**, E50, H38, T144, D139, E388, D392**;** Q33, S36, T37, R217, **Y87**, Q389, **S48**, T27, N29, R42, **Y61** | Nqo4 (E) |
|  | H60, E4, D49, E53; **S46**, **N48** | Nqo7 (P) |
|  | E223, E225, D220, E227, D303, D71, K240 E35, H233, E248**;** R216, Q226, R154, S158, R301, Y302, S139, S143, S145, K146, Y147, S148, S152, E235, R299, Y236, S237, S155, S156, Y232, T234, Q245, Y249, R294, **Q304**, **R307** | Nqo8 (Q) |

**Table SI.3. Average protonation microstates in ten loaded and ten unloaded snapshots**

| Type | Five E-Channel subunits | | Central-cluster residues | | N-side cluster residues | |
| --- | --- | --- | --- | --- | --- | --- |
|  | Loaded | Unloaded | Loaded | Unloaded | Loaded | Unloaded |
| Count Asp, Glu, His, Tyr, Arg, Lys | 224 | 224 | 15 | 15 | 30 | 30 |
| Residues that change | 74 | 74 | 6 | 6 | 5 | 5 |
| Number of unique protonation microstates | 26,360±1054 | 26,654±1102 | 32±8 | 34±8 | 18±7 | 24±7 |
| Unique protonation microstates in 90% of ensemble | 10,450±358 | 10,850±455 | 5±1 | 4±1 | 2±0 | 3±0 |
| Unique protonation microstates in 50% of ensemble | 236±60 | 240±45 | 3±0 | 3±0 | 1±0 | 1±0 |
| Average Charge | -3.14±0.3 | -3.88±0.3 | -1.10±0.1 | -1.99±0.2 | -2.87±0.2 | -3.94±0.3 |

Snapshots are classified by the average charge of the Central Cluster. Table 1 has statistics for a partially loaded snapshot.

**Table SI.4. Highest probability protonation microstates found in MCCE analysis of snapshots where the Central Cluster is loaded (top), mixed loaded-unloaded (middle) and unloaded (bottom).** The microstates are shown horizontally from the N to the P side (shapes: diamond: Glu, oval: Asp, triangle: His) Dark red: Glu^-^ or Asp^-^; dark blue: His^+^; Gray: neutral. The global average is the average of the probabilities of a microstate type found for different snapshots in MCCE simulation. The snapshots number are randomly selected representative snapshots.

| Loaded snapshots | | | | | | |
| --- | --- | --- | --- | --- | --- | --- |
| Microstates type | Total Charge | Probability in different snapshots | | | |  |
|  |  | Snapshot87 | Snapshot67 | Snapshot75 | Snapshot85 | Global Average |
| N-side P-side  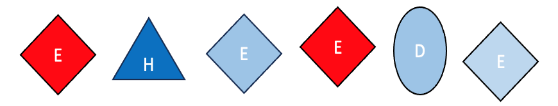 | -1 | 0.8% | 0.2% | 0.2% | 0.5% | 0.4% |
| 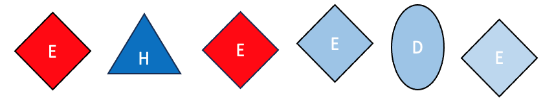 | -1 | 0.0% | 0.2% | 46.8% | 8.8% | 19.3% |
| 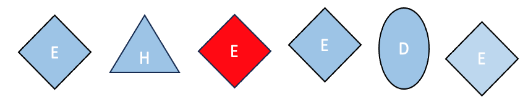 | -1 | 0.1% | 23.7% | 6.6% | 0.1% | 7.6% |
| 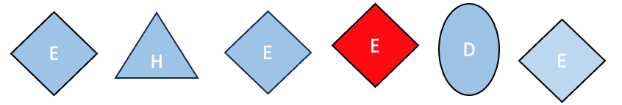 | -1 | 0.1% | 23.7% | 6.6% | 0.1% | 7.6% |
| 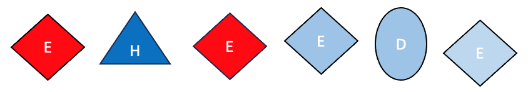 | -2 | 0.1% | 1.7% | 8.8% | 8.8% | 4.9% |
| 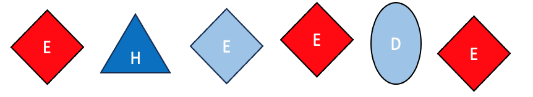 | -2 | 2.3% | 11.8% | 1.0% | 0.3% | 3.9% |
| Mix loaded-unloaded snapshots | | | | | | |
| Microstates type | Total Charge | Probability in different snapshots | | | |  |
|  |  | Snapshot61 | Snapshot64 | Snapshot76 | Snapshot85 | Global Average |
| N-side P-side  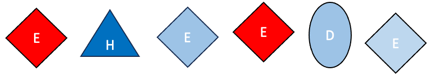 | -1 | 6.8% | 27.5% | 5.9% | 41.1% | 20.3% |
| 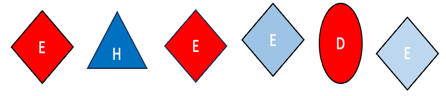 | -2 | 27.5% | 0.3% | 4.9% | 3.7% | 9.1% |
| 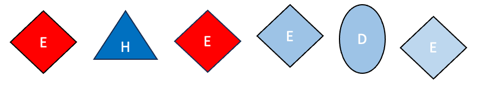 | -1 | 13.7% | 7.0% | 41.0% | 1.0% | 15.7% |
| 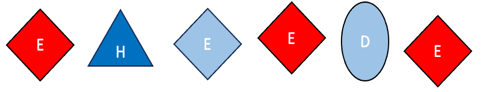 | -2 | 0.2% | 23.4% | 2.6% | 11.8% | 9.5% |
| 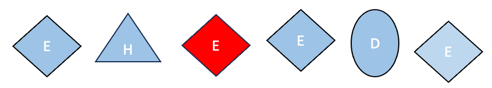 | -1 | 6.8% | 1.6% | 0.7% | 0.0% | 2.3% |
| 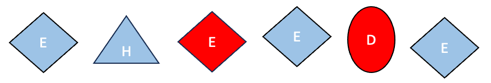 | -2 | 14.5% | 0.0% | 0.1% | 0.0% | 14.6% |
| 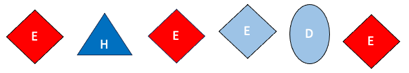 | -2 | 0.2% | 23.4% | 2.6% | 11.8% | 9.5% |
| Unloaded snapshots | | | | | | |
| Microstates type | Total Charge | Probability in different snapshots | | | |  |
|  |  | Snapshot0 | Snapshot65 | Snapshot71 | Snapshot86 | Global Average |
| N-side P-side  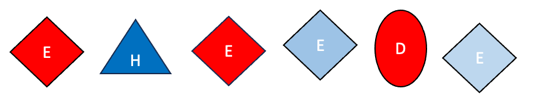 | -2 | 2.7% | 51.0% | 23.4% | 46.5% | 43.1% |
| 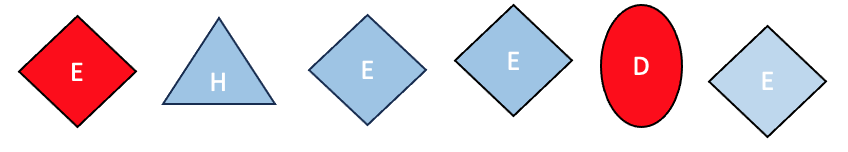 | -2 | 78.9% | 0.3% | 0.1% | 3.6% | 23.3% |
| 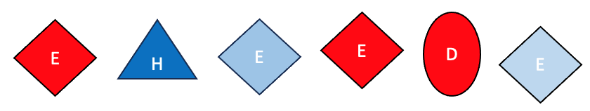 | -2 | 3.8% | 0.8% | 29.1% | 0.3% | 10.6% |
| 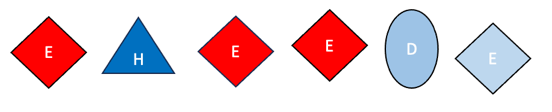 | -2 | 0.1% | 17.3% | 36.5% | 26.4% | 4.0% |
| 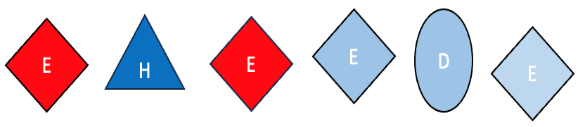 | -1 | 0.0% | 4.5% | 0.3% | 4.8% | 5.5% |
| 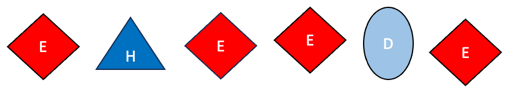 | -3 | 0.1% | 2.6% | 0.4% | 0.1% | 4.2% |

**Table SI.5. Experimental site-directed mutagenesis studies of the PLS residues in the N and Central Clusters and their conservation in Multiple Sequence Alignment.**

| Cluster | PLS (*T. thermophilus*) | Mutations, organism | Quinone activity | NADH activity | Ref. | MSA  Residue: % |
| --- | --- | --- | --- | --- | --- | --- |
| Central Cluster | 8-E163 | E157A, Ec | 24% |  | [33] | E:99.6; S:0.4 |
|  | 8-E213 | E212Q, Pd | 66% |  | [33] | E:99.6; A:0.4 |
|  | 7-D72 | D79N, Ec |  | 44% | [34] | D:100 |
|  | 7-E74 | E81A, Ec |  | 42% | [34] | E:100 |
|  | 7-D72/7-E74 | D79N/  E81Q, Ec |  | 2% | [34] | D:100/ E:100 |
|  | 8-H251 | None |  |  |  | H:44.8; A1.6; N: 51.6; S:1.6; G:0.4 |
|  | 8-E130 | None | ND | ND |  | E:44.8; S:54.3; G:0.8 |
| N-side cluster | 4-H38 | H228A, Ec | 33% |  | [35] | H:99.8; N:0.1 |
|  | 8-E225 | E216A, Ec | 80% |  | [36] | E:100 |
|  | 4-E51 | E36A |  | 20% | [36] | E:98.01; Y: 1.02 |
|  | 4-H58 | E71A |  | 48% | [36] | H:40.8; D:44.9; Q:2.8; R:0.3; Y:0.3; I:1.6; V:8.3; E:0.7; A:0.2 |
|  | 7-H60 | None | ND | ND |  | H:18.6; K:13.8; R:67.1; S:0.4; Q:0.1 |
|  | 4-E216 | None | ND | ND |  | E:33.5; Q: 9.5; K: 8.7; S:1.8; D:34.8; G:1.1; N:2.3 A:2.2; R:4.0; T:0.3; M:0.7; H:0.9; L:0.1; I:0.1 |
|  | 4-E50 | None | ND | ND |  | E:99.9 |

Complex I from *E. coli* (Ec), *P. denitrificans* (Pd), or *Y. lipolytica* (Yl). Activity refers to the measured NADH or quinone dependent activity relative to the wild type of enzyme. MSA is the result of multiple sequence alignment of 1000 Complex I sequences. See SI.1.6 for description of MSA.

**Table SI.6. List of the residues with the residues of within 5.0 Å of the active residues in the Central Cluster and the N-side cluster. Here, we considered the residues in the same chain for MSA analysis.**

| **Central Cluster Active residues** | **Residues within 5.0 Å volume of sphere of the Central Cluster active residues** |
| --- | --- |
| 8-E163 | **8-L159, 8-I160, 8-S161, 8-Y162, 8-L164, 8-G165, 8-L166, 8-E213** |
| 8-E213 | **8-L159, 8-S210, 8-M211, 8-A212, 8-A214, 8-A215, 8-L221** |
| 7-D72 | **7-F68, 7-F71, 7-V73** |
| 7-E74 | **7-L70, 7-F71, 7-D72, 7-V73, 7-V75, 7-A76, 7-F77, 7-L78, 7-W79, 7-F99, 7-L103** |
| 8-H251 | **8-A247, 8-G248, 8-Y249, 8-I250, 8-H251, 8-F252, 8-I253, 8-T254** |
| 8-E130 | **8-F126, 8-A127, 8-V128, 8-S129, 8-L131, 8-A132, 8-V133, 8-E163** |
| **N-side cluster Active residues** | **Residues within 5.0 Å volume of sphere of the N-side cluster active residues** |
| 4-H38 | **4-S36, 4-T37, 4-H38, 4-G39, 4-V40** |
| 8-E225 | **8-E223, 8-A224, 8-E227, 8-R299** |
| 4-E51 | **4-L47, 4-G49, 4-E50, 4-V52, 4-G387, 4-E388, 4-Q389, 4-V390** |
| 4-H58 | **4-L41, 4-R42, 4-V56, 4-P57, 4-H58, 4-I59** |
| 7-H60 | **7-P58, 7-V59, 7-F61, 7-Y62, 7-V63, 7-V115** |
| 4-E216 | **4-I213, 4-F214, 4-Y215, 4-R217, 4-A218** |
| 4-E50 | **4-L47, 4-G49, 4-E51, 4-Q389, 4-V390** |

The surrounding residues in the Central Cluster and the N-side cluster active residues **within 5.0 Å** of the volume sphere. These are in subunits Nqo4, Nqo7, and Nqo8 in *T. thermophilus* Complex I. Residues name identified by *T. thermophilus* subunits Nqo4-, Nqo7-, and Nqo8- one-letter residue name, and followed by residue ID.

Reference

[1]	Gupta, C., U. Khaniya, C.K. Chan, F. Dehez, M. Shekhar, M. Gunner, L. Sazanov, C. Chipot, and A. Singharoy, Charge transfer and chemo-mechanical coupling in respiratory complex I*.* Journal of the American Chemical Society, 2020. **142**(20): p. 9220-9230, <https://doi.org/10.1021/jacs.9b13450>.

[2]	Khaniya, U., C. Gupta, X. Cai, J. Mao, D. Kaur, Y. Zhang, A. Singharoy, and M. Gunner, Hydrogen bond network analysis reveals the pathway for the proton transfer in the E-channel of T. thermophilus Complex I*.* Biochimica et Biophysica Acta (BBA)-Bioenergetics, 2020. **1861**(10): p. 148240, <https://doi.org/10.1016/j.bbabio.2020.148240>.

[3]	Klauda, J.B., R.M. Venable, J.A. Freites, J.W. O’Connor, D.J. Tobias, C. Mondragon-Ramirez, I. Vorobyov, A.D. MacKerell Jr, and R.W. Pastor, Update of the CHARMM all-atom additive force field for lipids: validation on six lipid types*.* The journal of physical chemistry B, 2010. **114**(23): p. 7830-7843, <https://doi.org/10.1021/jp101759q>.

[4]	Best, R.B., X. Zhu, J. Shim, P.E. Lopes, J. Mittal, M. Feig, and A.D. MacKerell Jr, Optimization of the additive CHARMM all-atom protein force field targeting improved sampling of the backbone ϕ, ψ and side-chain χ1 and χ2 dihedral angles*.* Journal of chemical theory and computation, 2012. **8**(9): p. 3257-3273, <https://doi.org/10.1021/ct300400x>.

[5]	Chang, C.H. and K. Kim, Density functional theory calculation of bonding and charge parameters for molecular dynamics studies on [FeFe] hydrogenases*.* Journal of chemical theory and computation, 2009. **5**(4): p. 1137-1145, <https://doi.org/10.1021/ct800342w>.

[6]	Verkhovskaya, M.L., N. Belevich, L. Euro, M. Wikström, and M.I. Verkhovsky, Real-time electron transfer in respiratory complex I*.* Proceedings of the National Academy of Sciences, 2008. **105**(10): p. 3763-3767, <https://doi.org/10.1073/pnas.0711249105>.

[7]	Bridges, H.R., E. Bill, and J. Hirst, Mossbauer spectroscopy on respiratory complex I: the iron–sulfur cluster ensemble in the NADH-reduced enzyme is partially oxidized*.* Biochemistry, 2012. **51**(1): p. 149-158, <https://doi.org/10.1021/bi201644x>.

[8]	Roessler, M.M., M.S. King, A.J. Robinson, F.A. Armstrong, J. Harmer, and J. Hirst, Direct assignment of EPR spectra to structurally defined iron-sulfur clusters in complex I by double electron–electron resonance*.* Proceedings of the National Academy of Sciences, 2010. **107**(5): p. 1930-1935, <https://doi.org/10.1073/pnas.0908050107>.

[9]	Freddolino, P.L., K.H. Gardner, and K. Schulten, Signaling mechanisms of LOV domains: new insights from molecular dynamics studies*.* Photochemical & Photobiological Sciences, 2013. **12**(7): p. 1158-1170, <https://doi.org/10.1039/c3pp25400c>.

[10]	Song, Y., J. Mao, and M.R. Gunner, MCCE2: improving protein pKa calculations with extensive side chain rotamer sampling*.* Journal of computational chemistry, 2009. **30**(14): p. 2231-2247, <https://doi.org/10.1002/jcc.21222>.

[11]	Ranepura, G.A., J. Mao, J.V. Vermaas, J. Wang, C.J. Gisriel, R.J. Wei, J. Ortiz-Soto, M.R. Uddin, M. Amin, and G.W. Brudvig, Computing the Relative Affinity of Chlorophylls a and b to Light-Harvesting Complex II*.* The Journal of Physical Chemistry B, 2023. **127**(51): p. 10974-10986, <https://doi.org/10.1021/acs.jpcb.3c06273>.

[12]	Levy, R.M., L.Y. Zhang, E. Gallicchio, and A.K. Felts, On the nonpolar hydration free energy of proteins: surface area and continuum solvent models for the solute− solvent interaction energy*.* Journal of the American Chemical Society, 2003. **125**(31): p. 9523-9530, <https://doi.org/10.1021/ja029833a>.

[13]	Gunner, M., X. Zhu, and M.C. Klein, MCCE analysis of the pKas of introduced buried acids and bases in staphylococcal nuclease*.* Proteins: Structure, Function, and Bioinformatics, 2011. **79**(12): p. 3306-3319, <https://doi.org/10.1002/prot.23124>.

[14]	Zheng, Z. and M. Gunner, Analysis of the electrochemistry of hemes with Ems spanning 800 mV*.* Proteins: Structure, Function, and Bioinformatics, 2009. **75**(3): p. 719-734, <https://doi.org/10.1002/prot.22282>.

[15]	Baker, N.A., Improving implicit solvent simulations: a Poisson-centric view*.* Current opinion in structural biology, 2005. **15**(2): p. 137-143, <https://doi.org/10.1016/j.sbi.2005.02.001>.

[16]	Warwicker, J. and H. Watson, Calculation of the electric potential in the active site cleft due to α-helix dipoles*.* Journal of molecular biology, 1982. **157**(4): p. 671-679, <https://doi.org/10.1016/0022-2836(82)90505-8>.

[17]	Rocchia, W., E. Alexov, and B. Honig, Extending the applicability of the nonlinear Poisson− Boltzmann equation: multiple dielectric constants and multivalent ions*.* The Journal of Physical Chemistry B, 2001. **105**(28): p. 6507-6514, <https://doi.org/10.1021/jp010454y>.

[18]	Sitkoff, D., K.A. Sharp, and B. Honig, Accurate calculation of hydration free energies using macroscopic solvent models*.* The Journal of Physical Chemistry, 1994. **98**(7): p. 1978-1988, <https://doi.org/10.1021/j100058a043>.

[19]	Cornell, W.D., P. Cieplak, C.I. Bayly, I.R. Gould, K.M. Merz, D.M. Ferguson, D.C. Spellmeyer, T. Fox, J.W. Caldwell, and P.A. Kollman, A second generation force field for the simulation of proteins, nucleic acids, and organic molecules*.* Journal of the American Chemical Society, 1995. **117**(19): p. 5179-5197, <https://doi.org/10.1021/ja00124a002>.

[20]	Beroza, P., D. Fredkin, M. Okamura, and G. Feher, Protonation of interacting residues in a protein by a Monte Carlo method: application to lysozyme and the photosynthetic reaction center of Rhodobacter sphaeroides*.* Proceedings of the National Academy of Sciences, 1991. **88**(13): p. 5804-5808, <https://doi.org/10.1073/pnas.88.13.5804>.

[21]	Yang, A.S., M. Gunner, R. Sampogna, K. Sharp, and B. Honig, On the calculation of pKas in proteins*.* Proteins: Structure, Function, and Bioinformatics, 1993. **15**(3): p. 252-265, <https://doi.org/10.1002/prot.340150304>.

[22]	Wei, R.J., U. Khaniya, J. Mao, J. Liu, V.S. Batista, and M. Gunner, Tools for analyzing protonation states and for tracing proton transfer pathways with examples from the Rb. sphaeroides photosynthetic reaction centers*.* Photosynthesis research, 2023. **156**(1): p. 101-112, <https://doi.org/10.1007/s11120-022-00973-0>.

[23]	Zhang, Y., K. Haider, D. Kaur, V.A. Ngo, X. Cai, J. Mao, U. Khaniya, X. Zhu, S. Noskov, and T. Lazaridis, Characterizing the water wire in the Gramicidin channel found by Monte Carlo sampling using continuum electrostatics and in molecular dynamics trajectories with conventional or polarizable force fields*.* Journal of Computational Biophysics and Chemistry, 2021. **20**(02): p. 111-130, <https://doi.org/10.1142/S2737416520420016>.

[24]	Liu, Y., Q. Meng, R. Chen, J. Wang, S. Jiang, and Y. Hu, A new method to evaluate the similarity of chromatographic fingerprints: weighted pearson product-moment correlation coefficient*.* Journal of chromatographic science, 2004. **42**(10): p. 545-550, <https://doi.org/10.1093/chromsci/42.10.545>.

[25]	Sievers, F., A. Wilm, D. Dineen, T.J. Gibson, K. Karplus, W. Li, R. Lopez, H. McWilliam, M. Remmert, and J. Söding, Fast, scalable generation of high‐quality protein multiple sequence alignments using Clustal Omega*.* Molecular systems biology, 2011. **7**(1): p. 539, <https://doi.org/10.1038/msb.2011.75>.

[26]	Crooks, G.E., G. Hon, J.-M. Chandonia, and S.E. Brenner, WebLogo: a sequence logo generator*.* Genome research, 2004. **14**(6): p. 1188-1190, <https://doi.org/10.1101/gr.849004>.

[27]	Kaila, V.R., Resolving chemical dynamics in biological energy conversion: Long-range proton-coupled electron transfer in respiratory complex I*.* Accounts of Chemical Research, 2021. **54**(24): p. 4462-4473, <https://doi.org/10.1021/acs.accounts.1c00524>.

[28]	Zheng, W., P. Chai, J. Zhu, and K. Zhang, High-resolution in situ structures of mammalian respiratory supercomplexes*.* Nature, 2024: p. 1-8, <https://doi.org/10.1038/s41586-024-07488-9>.

[29]	Gunner, M., J. Madeo, and Z. Zhu, Modification of quinone electrochemistry by the proteins in the biological electron transfer chains: examples from photosynthetic reaction centers*.* Journal of bioenergetics and biomembranes, 2008. **40**: p. 509-519, <https://doi.org/10.1007/s10863-008-9179-1>.

[30]	Sazanov, L.A., A giant molecular proton pump: structure and mechanism of respiratory complex I*.* Nature reviews Molecular cell biology, 2015. **16**(6): p. 375-388, <https://doi.org/10.1038/nrm3997>.

[31]	Wirth, C., U. Brandt, C. Hunte, and V. Zickermann, Structure and function of mitochondrial complex I*.* Biochimica et Biophysica Acta (BBA)-Bioenergetics, 2016. **1857**(7): p. 902-914, <https://doi.org/10.1016/j.bbabio.2016.02.013>.

[32]	Grba, D.N. and J. Hirst, Mitochondrial complex I structure reveals ordered water molecules for catalysis and proton translocation*.* Nature structural & molecular biology, 2020. **27**(10): p. 892-900, <https://doi.org/10.1038/s41594-020-0473-x>.

[33]	Kurki, S., V. Zickermann, M. Kervinen, I. Hassinen, and M. Finel, Mutagenesis of three conserved Glu residues in a bacterial homologue of the ND1 subunit of complex I affects ubiquinone reduction kinetics but not inhibition by dicyclohexylcarbodiimide*.* Biochemistry, 2000. **39**(44): p. 13496-13502, <https://doi.org/10.1021/bi001134s>.

[34]	Kao, M.-C., S. Di Bernardo, M. Perego, E. Nakamaru-Ogiso, A. Matsuno-Yagi, and T. Yagi, Functional roles of four conserved charged residues in the membrane domain subunit NuoA of the proton-translocating NADH-quinone oxidoreductase from Escherichia coli*.* Journal of Biological Chemistry, 2004. **279**(31): p. 32360-32366, <https://doi.org/10.1074/jbc.M403885200>.

[35]	Belevich, G., L. Euro, M. Wikström, and M. Verkhovskaya, Role of the conserved arginine 274 and histidine 224 and 228 residues in the NuoCD subunit of complex I from Escherichia coli*.* Biochemistry, 2007. **46**(2): p. 526-533, <https://doi.org/10.1021/bi062062t>.

[36]	Sinha, P.K., J. Torres-Bacete, E. Nakamaru-Ogiso, N. Castro-Guerrero, A. Matsuno-Yagi, and T. Yagi, Critical roles of subunit NuoH (ND1) in the assembly of peripheral subunits with the membrane domain of Escherichia coli NDH-1*.* Journal of biological chemistry, 2009. **284**(15): p. 9814-9823, <https://doi.org/10.1074/jbc.M809468200>.
